# Structural modeling of the hERG potassium channel and associated drug interactions

**DOI:** 10.1101/2021.07.08.451699

**Authors:** Jan Maly, Aiyana M. Emigh, Kevin R. DeMarco, Kazuharu Furutani, Jon T. Sack, Colleen E. Clancy, Igor Vorobyov, Vladimir Yarov-Yarovoy

## Abstract

The voltage-gated potassium channel, KV11.1, encoded by the human *Ether-à-go-go*- Related Gene (hERG) is expressed in cardiac myocytes, where it is crucial for the membrane repolarization of the action potential. Gating of hERG channel is characterized by rapid, voltage-dependent, C-type inactivation, which blocks ion conduction and is suggested to involve constriction of the selectivity filter. Mutations S620T and S641A/T within the selectivity filter region of hERG have been shown to alter the voltage-dependence of channel inactivation. Because hERG channel blockade is implicated in drug-induced arrhythmias associated with both the open and inactivated states, we used Rosetta to simulate effects of hERG S620T and S641A/T mutations to elucidate conformational changes associated with hERG channel inactivation and differences in drug binding between the two states. Rosetta modeling of the S641A fast-inactivating mutation revealed a lateral shift of F627 side chain in the selectivity filter into the central channel axis along the ion conduction pathway and formation of four lateral fenestrations in the pore. Rosetta modeling of the non-inactivating mutations S620T and S641T suggested a potential molecular mechanism preventing F627 side chain from shifting into the ion conduction pathway during the proposed inactivation process. Furthermore, we used Rosetta docking to explore the binding mechanism of highly selective and potent hERG blockers - dofetilide, terfenadine, and E4031. Our structural modeling correlate well with existing experimental evidence involving interactions of these drugs with key hERG residues Y652 and F656 inside the pore and reveal potential ligand binding interactions within fenestration region in an inactivated state.

**Significance Statement:** Computational models of hERG potassium channel provide structural insights into an inactivated state and associated drug interactions. Our computational approach will be useful to study ion channel modulation by small molecules.

## Introduction

The voltage-gated potassium channel, KV11.1, is encoded by the **h**uman ***E****ther-à-go-go*- **R**elated **G**ene (hERG) and plays a key role in cardiac myocytes, where it conducts the rapid delayed rectifier K^+^ current (*I*Kr) that regulates the repolarization phase of the ventricular action potential (AP), effectively controlling the duration of the heart’s QT interval on the surface electrocardiogram (ECG) (Smith et al., 1996; Vandenberg et al., 2012). This repolarization is shaped by the hERG channel’s unique C-type inactivation kinetics during membrane depolarization phase of the AP where, upon channel opening, the selectivity filter (SF) rapidly becomes constricted and abolishes the K^+^ conduction. This is followed by rapid recovery from inactivation and slow deactivation during membrane repolarization phase of the AP, thereby facilitating outward K^+^ current over the course of cardiac repolarization. Prolongation of the QT interval, also known as Long QT syndrome (LQTS), can be a consequence of drug-induced block of the hERG channel, and a reason for the withdrawal of multiple drugs from the pharmaceutical market and drug candidates from preclinical screenings (Curran et al., 1995; Fermini and Fossa, 2003; Hancox et al., 2008; Roden, 2004; Shah, 2002). A diverse set of drug molecules, from Class-III antiarrhythmics to cocaine (Guo et al., 2006), exhibit promiscuous binding to the hERG channel pore. Binding affinities of many hERG blocking drugs exhibit dependence on the channel’s conformational state (Perrin et al., 2008; Vandenberg et al., 2012). While some drugs, such as moxifloxacin and nifekalant, preferentially bind to the open hERG channel state and blocking conduction without pro-arrhythmic effects, others like dofetilide and d-sotalol exhibit a higher affinity to the channel inactivated state and suggested to increase their pro-arrhythmic tendencies (Alexandrou et al., 2006; Ficker et al., 2001; Kushida et al., 2002; Numaguchi et al., 2000; Romero et al., 2015). Thus, the mechanisms by which many structurally diverse compounds preferentially block hERG channels in a state-dependent manner are of significant interest for drug safety (Perrin et al., 2008; Wu et al., 2015).

The crucial role of hERG channel in cardiac AP repolarization and its susceptibility to a vast array of drugs are reliant on its unique structural features. Like other voltage-gated K^+^ (Kv) channels, hERG is comprised of six transmembrane segments (S1-S6), where S1-S4 form the voltage-sensing domain (VSD) and S5-S6 with the intervening extracellular turret (S5P) and pore helix (P-helix) making up the pore-forming domain. The majority of KV channels have a domain-swapped voltage sensor arrangement, whereby conformational changes are transmitted from the VSD to the pore domain by a 15 residue-long helical S4-S5 linker that interacts with the S6 segment from adjacent subunit (Long et al., 2005a; Long et al., 2005b; Lu et al., 2001). However, in the cryo-electron microscopy (cryoEM) structure of hERG, the linker between the S4 and S5 segments is only 4-residue-long, suggesting a more direct coupling of conformational change from the VSD to pore domain of the same subunit during channel gating and suggesting a possible role in hERG’s rapid and voltage-dependent inactivation (Fernandez-Marino et al., 2018; Schonherr and Heinemann, 1996; Smith et al., 1996; Spector et al., 1996b; Wang and MacKinnon, 2017). The long S5P linker (about 40 residues in hERG) is significantly different between hERG and other KV channels and presents another possible link to hERG inactivation. Charge reversal mutagenesis and domain-swapping between hERG and the closely-related EAG1 channel suggests a close link between the turret linker and rapid C-type inactivation (Brugada et al., 2004; Clarke et al., 2006; Herzberg et al., 1998). Lastly, the pore cavity of hERG presents a unique feature found in the cryoEM structure that sets it apart from other KV channels – hydrophobic pockets extending from the central cavity just below the SF to a narrow space between the S6 and P helices, with a depth of ∼11 Å, and potential to accommodate parts of hERG-blocking drugs (Wang and MacKinnon, 2017). In addition to these hydrophobic pockets, the S6 segments and P-helices form a relatively narrow central cavity with a strong electronegative potential below the SF, allowing for a potentially tighter mode of binding for many hERG blockers upon gate opening (Wang and MacKinnon, 2017).

Another important feature for drug binding and efficacy is hERG inactivation. Like other KV channels, K^+^ flux through the hERG channel is regulated by the processes of activation, deactivation and inactivation. Activation initiates K^+^ flux through the pore via the opening of its cytoplasmic gate in response to membrane depolarization whereas channel pore closure, or deactivation, occurs during membrane repolarization. Lastly, inactivation stops the flow of K^+^ ions through the pore at depolarized membrane potentials. Initial reports of C-type inactivation in the Shaker channel described it as a slower and voltage-independent form of inactivation, sensitive to external tetraethylammonium (TEA) ion concentration and insensitive to cytoplasmic N-terminus deletion (critical to so-called “ball and chain” N-type inactivation), suggesting that C-type inactivation blocks ion flow through conformational changes at the SF (Choi et al., 1991; Hoshi et al., 1991; Zhou et al., 2001a). Furthermore, a crystal structure of the KcsA channel identified four SF K^+^ binding sites, S1 to S4, formed by backbone carbonyl oxygen atoms that line up and protrude into the conduction pathway of the SF, from the extracellular to intracellular side (Zhou et al., 2001b). Analysis of the KcsA crystal structure showed that K^+^ ions are coordinated by the carbonyl oxygens of the SF residues, that conduction proceeds through a staggered occupancy of the SF sites by water and K^+^ ions, and that at low K^+^ concentrations, the filter loses coordination of K^+^ ions at the S2-S3 position, inducing a conformational change akin to “collapse” or “constriction” that renders it non-conductive (Zhou et al., 2001a; Zhou et al., 2001b). Further support for this form of C-type inactivation was demonstrated by an inactivated state KcsA crystal structure that, while sharing similar conformational consequences such as de-coordination of K^+^ ions at the S2-S3 position and SF constriction, suggested that the extent of inner gate opening (TM2 helix in KcsA; S6 helix in KV), rather than low extracellular K^+^ concentration, was the key allosteric mechanism involving conformational changes within the SF resulting in a C-type inactivated state (Cuello et al., 2010a; Cuello et al., 2010b). Alternatively, a recent crystal structure of a KV1.2-2.1 chimeric channel bearing an inactivation-inducing mutation supports the notion that a low external K^+^ concentration is a major contributing factor to C-type inactivation, since de-coordination of a top SF K^+^ ion due to a slight reorientation of the backbone carbonyl at the S1 position is enough to block K^+^ ion conduction (Pau et al., 2017). Similar to the Shaker channel, C-type inactivation in hERG channel is sensitive to external application of TEA and ion occupancy of the SF, with increasing K^+^ ion concentrations attenuating inactivation (Lopez-Barneo et al., 1993; Schonherr and Heinemann, 1996; Smith et al., 1996). In contrast to Shaker, however, the rate of hERG inactivation is fast and voltage-dependent (Schonherr and Heinemann, 1996; Wang et al., 1997). While there is consensus that the SF is the site of C-type inactivation and that structural rearrangements involving de-coordination of K^+^ ions disrupt ion flux, the exact nature of these conformational changes remains unclear, and is most likely tied to unique structural differences between hERG and other KV channels.

The ability of drugs to enter the hERG cavity allows them to access many features of the hERG structure. Located inside the pore, and lining the K^+^ conduction pathway, are residues that are important molecular determinants for high-affinity binding of many hERG blocking drugs, including MK-499, cisapride, dofetilide, terfenadine, and E-4031 (Kamiya et al., 2006; Kamiya et al., 2008; Mitcheson et al., 2000). Key residues forming hERG drug receptor sites include Y652 and F656, aromatic residues located on each of the four S6 segments within the central cavity and critical for high-affinity interactions with a variety of structurally diverse compounds, as well as residues T623, S624 and V625 at the bottom of the SF and P-helix (Fernandez et al., 2004; Mitcheson et al., 2000; Mitcheson and Perry, 2003; Perry et al., 2010; Sanguinetti and Mitcheson, 2005; Vandenberg et al., 2012). Among the wide variety of cardiac and noncardiac drugs known to bind to the channel with the potential to cause arrhythmias, several drugs have been studied extensively due to their action as Class III antiarrhythmic agents or their inadvertent ability to act as potent hERG blockers. Dofetilide and E-4031 are Class III antiarrhythmic methane sulfonamides developed to preferentially block *I*Kr current and prolong cardiac action potential duration (Spector et al., 1996a). Both drugs share similar potency against the wild-type (WT) hERG channel, with IC50 in the low nanomolar range and significantly reduced channel block of hERG alanine mutants of Y652 and F656 and T623, S624, V625 (Kamiya et al., 2006; Orvos et al., 2019; Perrin et al., 2008; Zhou et al., 1998). Unlike dofetilide and E-4031, terfenadine represents an example of a noncardiac drug capable of inducing cardiac arrhythmias and Long QT syndrome. Terfenadine is an antihistamine formerly used to treat a variety of allergy symptoms but withdrawn from the pharmaceutical market due to cardiotoxicity at higher dosages or with prolonged use (Perrin et al., 2008). Notably, its direct metabolite fexofenadine contains a terminal carboxylic acid functional group and does not display any cardiotoxicity (Roy et al., 1996; Suessbrich et al., 1996). Similar to dofetilide and E-4031, mutations of hERG F656 and Y652 residues to alanine attenuate block by terfenadine, although the drug does not appear to be affected by alanine mutants of hERG SF residues (Mitcheson et al., 2000). The three drugs illustrate the variability in structure and function of drugs that block hERG, and a reason why all new pharmaceutical compounds must undergo screening for hERG/*I*Kr block during drug development (Gintant et al., 2006; Hancox et al., 2008).

Just as there are mutations directly affecting drug binding to the hERG channel, there are numerous mutations that modify hERG inactivation and consequently exert varied effects on drug binding. Early studies demonstrating the importance of the region surrounding the SF showed that residues S620 on the P-helix, S631 on the S6 segment-SF loop, and S641 near the top of the S6 segment were critical to fast C-type inactivation in the hERG channel (Vandenberg et al., 2012). Replacing S620 with threonine (S620T) has been shown to abolish inactivation and significantly attenuate dofetilide block of hERG, despite the presence of critical drug-block residues Y652/F656 (Ficker et al., 2001; Herzberg et al., 1998). Similarly, the S631A mutation exhibits strong attenuation of inactivation with an associated drop in dofetilide efficacy, even though S631 is located outside of the proposed drug-binding region (Ficker et al., 2001; Zou et al., 1998). Although the results of these studies may imply a direct correlation between inactivation and high-affinity drug binding in hERG, a recent study has shown that while a single hERG subunit harboring the S620T mutation was enough to completely abolish inactivation, drug potency of dofetilide, MK-499, and cisapride was directly proportional to the number of mutant subunits in the channel, unrelated to the level of inactivation; results for S631A, however, showed a direct correlation between both inactivation and dofetilide potency with increasing number of mutant subunits (Wu et al., 2015). Conversely, S641A mutation has been shown to strongly increase hERG channel inactivation (shifting it to more negative potentials), but also to attenuate binding of E-4031 (Bian et al., 2004). Other mutations such as N588K, located on the turret (S5P helix), G648A (S6 helix), F627Y (SF), and residues 623 – 625 on the pore helix all show varied effects on inactivation and hERG block by a multitude of different drug compounds, indicating a complex relationship between multiple allosteric conformational changes throughout the hERG channel that tie into the inactivation kinetics and drug-induced channel blockade (Butler et al., 2019; Clarke et al., 2006; Guo et al., 2006; Mitcheson et al., 2000).

In an effort to elucidate the structural determinants of inactivation and subsequent state-dependent drug binding of the hERG channel, we utilized existing structural data from the recent hERG cryoEM structure (Wang and MacKinnon, 2017) as a template for Rosetta modeling studies. To this end, we used *in silico* methods to incorporate inactivation-enhancing (S641A) and non-inactivating mutations (S641T/S620T) into hERG WT models. We used Rosetta relax protocols (Nivon et al., 2013) to observed conformational changes associated with the S641A inactivation-enhancing mutant, including an inward shift of F627 on the SF and formation of four lateral fenestrations in the pore near a hydrophobic patch of key drug-binding residues Y652 and F656 on S6, and F557 on S5. Additionally, we show that non-inactivating mutations S620T and S641T block the inward shift of F627 via differing mechanisms, preventing potential conformational changes of the SF during inactivation. Lastly, we used Rosetta GALigandDock protocol (Park et al., 2021) to explore interactions of dofetilide, terfenadine and E4031 with the WT and mutant hERG channel structural models. Our results show that while dofetilide, terfenadine and E4031 bound to key residues Y652 and F656 across all models, in agreement with existing experimental evidence, only the S641A inactivation-enhancing mutant models revealed alternative state-dependent drug binding.

## Materials and Methods

### WT structures and model selection

hERG channel WT models 1 (m1) and 2 (m2) were generated from the published putatively open-state hERG structure (Wang and MacKinnon, 2017) (PDBID: 5VA2). For m1, the original cryoEM structure was first passed through RosettaCM (Song et al., 2013) to rebuild missing loop regions and then refit using the Rosetta cryoEM refinement protocol (DiMaio et al., 2015). For m2, the original cryoEM structure was first refit using the Rosetta cryoEM refinement protocol and then passed through RosettaCM to rebuild missing loop regions. For the cryoEM refinement step, the lowest energy models were visually inspected before the lowest-scoring model was selected. For RosettaCM, the top 10% of 10,000 generated structures were extracted, and the lowest energy structure was selected for study.

### Mutant structures and model selection

All mutants were created from WT m1 and m2 models using amino acid replacement in UCSF Chimera and a membrane topology file generated for the WT using a combination of the hERG cryoEM structure and Octopus web server (Pettersen et al., 2004; Viklund and Elofsson, 2008). Using the two models as templates, Rosetta symmetry definition files were created for each model, and models were minimized using a single run of Rosetta FastRelax with symmetry, using the ref2015 (specific to Rosetta version 3.11, Rv3.11) or talaris2014 (specific to the older Rosetta version 3.6, Rv3.6) score function and membrane weights with Cartesian coordinates (membrane_highres_Menv_smooth_cart) (Alford et al., 2017; Conway et al., 2014; Nivon et al., 2013). With the exception of the S641A m1 models which were generated using Rosetta version 3.6, all models were updated to Rosetta version 3.11. Additionally, two types of movemap files were used to define which backbone and sidechain torsion angles are moveable in each run, with the radial (rad) movemap specifying residues within a roughly 15 A radius from the center of the SF, and the regional (reg) movemap specifying the entire pore domain (S5, S6, S5P, and P helices and the SF). The details of the movemap files and generalized command line markup are described in the Supplemental Material. We generated 10,000 hERG channel models in each round of Rosetta Relax, and the top 10% models by total score were extracted and RMSD from the WT cryoEM refined models was calculated. RMSD values were calculated for the whole-protein against input structures using Rosetta, and for specific amino acids using UCSF Chimera (Pettersen et al., 2004). UCSF Chimera was also used to calculate RMSD between top-scoring and input models using superposition of the P-helix, and to calculate residue angles consisting of three carbon atoms (typically Cα, C and Cψ), as well as torsion angles using Cα, C , Cψ, and C atoms. Aromatic centroids and distances between them were measured using Chimera. Residue-specific interaction energies were calculated using the Rosetta residue_energy_breakdown application from which *total*, *fa_attr*, *fa_rep*, *fa_sol*, *hbond_bb_sc*, and any other relevant interaction parameter scores were extracted. With the exception of the S641A Rv3.6 m1 models, the top-scoring Rv3.11 regional models from each run were used for subsequent docking simulations.

### Ligand docking

Structures of terfenadine (terf) and dofetilide (dft) in the mol2 format were downloaded from the ZINC15 compound database(Sterling and Irwin, 2015). Chemical structure of E- 4031 was drawn using ChemDraw (PerkinElmer) and converted to a 3D structure using OpenBabel (O’Boyle et al., 2011). For positively charged versions of each drug, hydrogens were added to the central nitrogen and a +1 charge was added using the Antechamber function in UCSF Chimera (Pettersen et al., 2004). Ligand energy minimization was performed using the “Optimize Geometry” function in Avogadro using the generalized AMBER force field (GAFF) (Hanwell et al., 2012). Ligand parameters files were generated using Rosetta scripts and incorporated in the Rosetta docking protocol. For docking ligands to the hERG channel models, we utilized the RosettaScripts GALigandDock (Park et al., 2021) method using the virtual screening, high accuracy mode and allowing for amino acid sidechain and backbone flexibility within the binding pocket and additional cartesian minimization of +1 sidechains up and downstream. Using Chimera, ligand files (mol2 or pdb) were placed within the pore lumen region just below the SF and saved as a single pdb file for input to GALigandDock. The generalized command line and XML markup are described in the Supplemental Material. Each GALigandDock simulation generated 2,000 models and the top 50 models were identified by Ligand Interface score. Protein-ligand interactions were evaluated using the Protein-Ligand Interaction server (PLIP) (Salentin et al., 2015).

## Results

### hERG WT input structures

To generate mutant channel models and explore associated conformational changes, WT input models with complete loop segments (those missing in the cryoEM structures) were developed first (see Methods). Since loop regions tend to be highly dynamic, we used two starting WT hERG channel structural models (hERG WT m1 and hERG WT m2) that differ primarily in their loop conformations to allow for larger structural sampling. Both WT hERG input models were generated using the cryoEM hERG channel structure (Wang and MacKinnon, 2017) and refined (with loop rebuilding) using Rosetta’s cryoEM refinement protocol (DiMaio et al., 2015). Both models exhibit near identical domain structure with voltage-sensing and pore domains (VSD and PD, respectively) organized in a non-domain-swapped homo-tetrameric arrangement (**Figure 1A, hERG WT m2 depicted**). In the SF, residues F627 are pointing away from the pore central axis, while S624 at the bottom of the SF are pointing into the pore central axis and pore lumen (**Figure 1A, top-view detail**). Along the S5 and S6 segments, residues Y652, F656, and F557 are tightly packed, with the aromatic groups of Y652 and F656 in a near-parallel arrangement, and those of F656 and F557 in a near-perpendicular arrangement, suggestive of stable π-π interactions (**Figure 1A, side-view detail**). While backbone conformation and positioning of the key residues is nearly identical for both structures, large structural deviations arise in the loop regions near the extracellular side of each segment, with conformational differences up to 16 Å between the S1 and S2 segments; a result of rebuilding variable loop regions of the cryoEM structure (**Figure 1B**). Of particular interest is the position of R582, a residue implicated in conformational changes of the S5P “turret” helix during inactivation (Clarke et al., 2006; Fougere et al., 2011). In the hERG WT m1 model, R582 is positioned away from the turret, pointing into the extracellular space (**Figure 1B**). In the hERG WT m2 model, R582 is positioned just above the SF, and within hydrogen bonding distance to N588 and D591 (**Figure 1B, Supplemental Fig. 1A inset**). As described below, the positioning of R582 may play a critical role in hERG conformational stability and inactivation, enabling formation of the fenestrations (not found in the WT models) between the S5, S6 and P segments just below the SF, and allowing dynamic movements of other key residues.

**Fig. 1.**
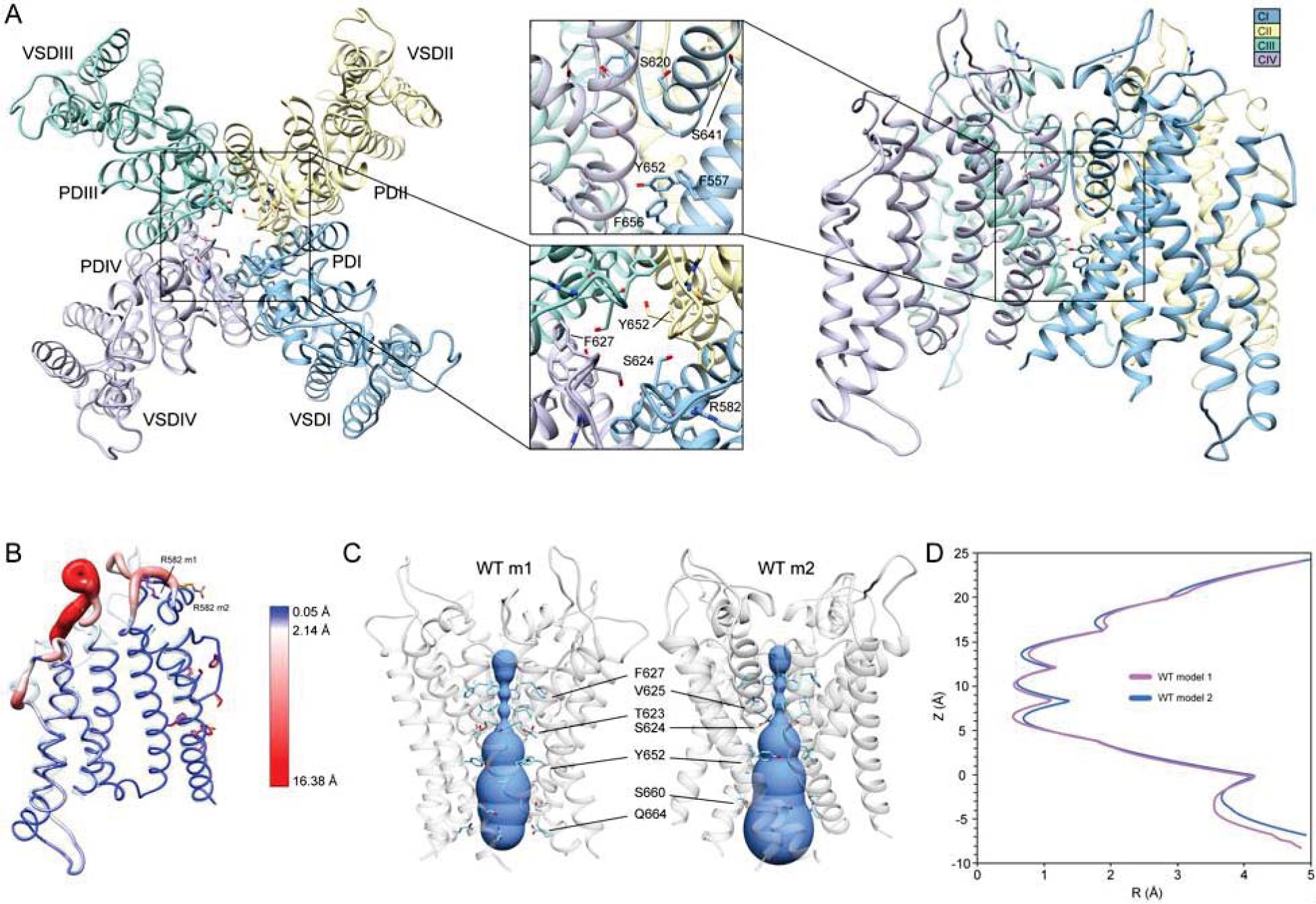
hERG WT channel model 1 (m1) and model 2 (m2) comparison. (A) Domain structure of hERG WT channel. Chains are colored according to the legend as CI-CIV. PD: pore domain, VSD: voltage sensing domain. (B) Overlay of hERG WT m1 and m2 models. M1 model is in translucent light-blue cartoon with purple residues, m2 model in noodle wireframe and color coded according to RMSD, with orange residues. (C) Van der Waals pore radius comparison for hERG WT m1 and m2 models. Volume spheres are in blue, with surrounding residues in light blue. (D) hERG WT m1 (purple) and m2 (blue) pore radius, *R*, profiles as a function of *z* generated by the HOLE program (Smart et al., 1996).

Another key difference between the two hERG WT models is the van der Waals pore radius (**Figure 1C, D**). Both models reveal constrictions at the K^+^/H2O-coordinating carbonyl oxygens in the GFG segment of the SF, with additional constrictions formed by Y652 and S660 in the middle of the pore lumen. The regions near the extracellular and intracellular gates however differ slightly in their pore radii. While the hERG m2 model shows an expanded pore volume at the intracellular gate, the hERG m1 model’s pore is significantly narrower, pinched by Q664 sidechains. The reverse is true near the extracellular gate at the top of the SF, where the pore radius appears to be smaller in the m2 model compared to m1. Given that the conformation of the turret helix is nearly identical in both models, this slight difference can be attributed to the positioning of the R582 sidechain, which dips down just above the SF, encircling the pore axis. Overall, the effect is a slightly narrower pore cavity and SF region in the hERG WT m1 model, and a slightly narrower extracellular gate region in hERG WT m2.

Conformations and Rosetta energy interactions of amino acids associated with the various mutations described herein were similar between hERG WT m1 and hERG WT m2 starting structures and are shown in **Supplemental Fig. 1**. The position of F627 is nearly identical in both models, with an all-atom RMSD of 0.42 Å; F627 in model 2 does however exhibit a stronger interaction with F617 of the neighboring chain (-1.30 kcal/mol compared to -0.77 kcal/mol in model 1), suggesting a more tightly packed pore domain structure. RMSDs of surrounding/contacting residues are equal to or less than that of F627 (**Supplemental Fig. 1A**). Additionally, the S620 sidechain oxygen atom is oriented away from the SF backbone in all chains of model 2, whereas it is pointed towards the SF backbone in two chains of model 1. While this orientation does increase the interaction energy between S620 and the G626 backbone due to hydrogen bonding (measured using Rosetta’s *hbond_bb_sc* energy term), the overall RMSD of the GFG motif of the SF is only about 0.4 Å (**Supplemental Fig. 1A**). Similarly, two chains of model 1 show S649 hydroxyl groups pointing away from the Y652 hydroxyl group of the adjacent chain, with the other two pointing towards them; all chains of model 2 show S649 hydrogen bonding with Y652 (using the *hbond_sc* energy term). While the S649 hydrogen bonds increase the overall interaction energy with Y652, they create a steric clash with F656, lowering the total interaction energy. Despite these minor differences, the overall all-atom and backbone RMSDs for S649, Y652, F656, and F557 are 0.2 to 0.3 Å, with the aromatic planes of F656 parallel to Y652 (with over -7.0 kcal/mol of interaction energy) and perpendicular to F557 (approximately -2 kcal/mol) (**Supplemental Fig. 1A**).

To explore differences in conformational stability between the WT m1 and m2 models, we used RosettaRelax to perturb each model through a series of repacking and minimization steps (see Methods). Results of RosettaRelax begin to illustrate some key differences between the hERG WT model 1 and 2 structures and subsequent mutations, specifically movement of F627 into the conduction path, conformational changes opening up the hydrophobic patch of F557, Y652, and F656, and orientation of S620 behind the SF. In relaxed WT m1 structures, both radial and regional relax models (those using a movemap encompassing the pore domain within a 15 Å radius from the center of the SF during the RosettaRelax protocol versus those that use the entire pore domain, respectively; see Methods) showed a 2.0 to 2.7 Å shift of F627 into the conduction path (distances measured between aromatic centroids) (**Supplemental Fig. 1B**). Furthermore, the hydrophobic patch along the S5 and S6 segments in the relaxed WT m1 regional models exhibit similar positioning of residues to the input structure, with F656 showing only slight rotation but remaining perpendicular to F557. Likewise, Y652 remains relatively parallel to F656, deviating from this position only when S649 on the adjacent chain shifts the hydrogen bond with Y652 upwards, along the conduction path; hydrogen bond interactions of about -2.8 kcal/mol between S649 and Y652 are present in all relaxed WT m1 and m2 structures (**Supplemental Fig. 1B**). Larger conformational changes in this region can be seen in m1 radial models, where F656 shifts further into the pore cavity, losing its interaction with F557. This shift appears to be greatly facilitated by the upward shift of S649 hydrogen bonding to Y652, which changes its orientation to F656 from parallel to perpendicular (**Supplemental Fig. 1B**).

Conformational changes in WT m2 models are significantly lower across all chains than in m1 models, preventing the large deviations between the input and relaxed models in the aforementioned residues. Compared to the input model, distances of F627 aromatic centroids across all relaxed m2 models are nearly identical, with only a 0.5 Å shift into the conduction path, a possible effect of the S620 hydroxyl oxygen hydrogen bonding with the F627 backbone NH group in all m2 models (**Supplemental Fig. 1C**). Furthermore, F656 retains a near perpendicular orientation to F557, with an interaction energy of approximately -2 kcal/mol despite a significant upward shift of the S649-Y652 hydrogen bonding pair position (**Supplemental Fig. 1C**). The much larger conformational freedom of m1 models begins to illustrate how the various mutations, described below, can affect structural changes between open and inactivated states of hERG.

### Drug docking to hERG WT

To determine the mode of drug binding to the WT models, we used Rosetta GALigandDock (Park et al., 2021) to dock dofetilide (dft), terfenadine (terf) and E4031 to top-scoring relaxed WT structures. Drug docking showed some distinct differences between the hERG WT m1 and m2 models. For docking of neutral and charged dofetilide (dft-0 and dft-1, respectively) to hERG WT m1, top-scoring models showed hydrophobic and aromatic π−π interactions with Y652 and A653 for both charged and neutral ligands, in addition to hydrogen bonding between the central and sulfonamide nitrogens (for dft-1) and sulfonamide oxygens (for dft-0) with Y652 hydroxyl groups (**Figure 2A**). For the m2 model, dft-0 showed similar hydrophobic interactions with A653 but was shifted further down the pore to interact with F656 and formed a hydrogen bond with backbone carbonyl of A653. A similar downward shift was observed with dft-1, with increased hydrogen bonding to backbone carbonyl of G657. This downward shift was seen in all m2-docked dofetilide models, with the majority of top-scoring models located in the mid-pore region or near the intracellular gate. The reverse was true for dofetilide docked to m1 models, where primary hydrophobic interactions involved Y652, and hydrogen bonding with S624 was observed for lower scoring charged ligands (within the top 50 scored structures). A possible explanation for this may lie in the increased conformational freedom of m1 models, allowing for a larger opening of, and therefore increased access to, the upper regions of the hERG cavity. Overall, top-scoring dofetilide structures maintained a bent orientation in both m1 and m2 models, maximizing exposure to hydrophobic and polar sidechain and backbone atoms along the pore.

**Fig. 2.**
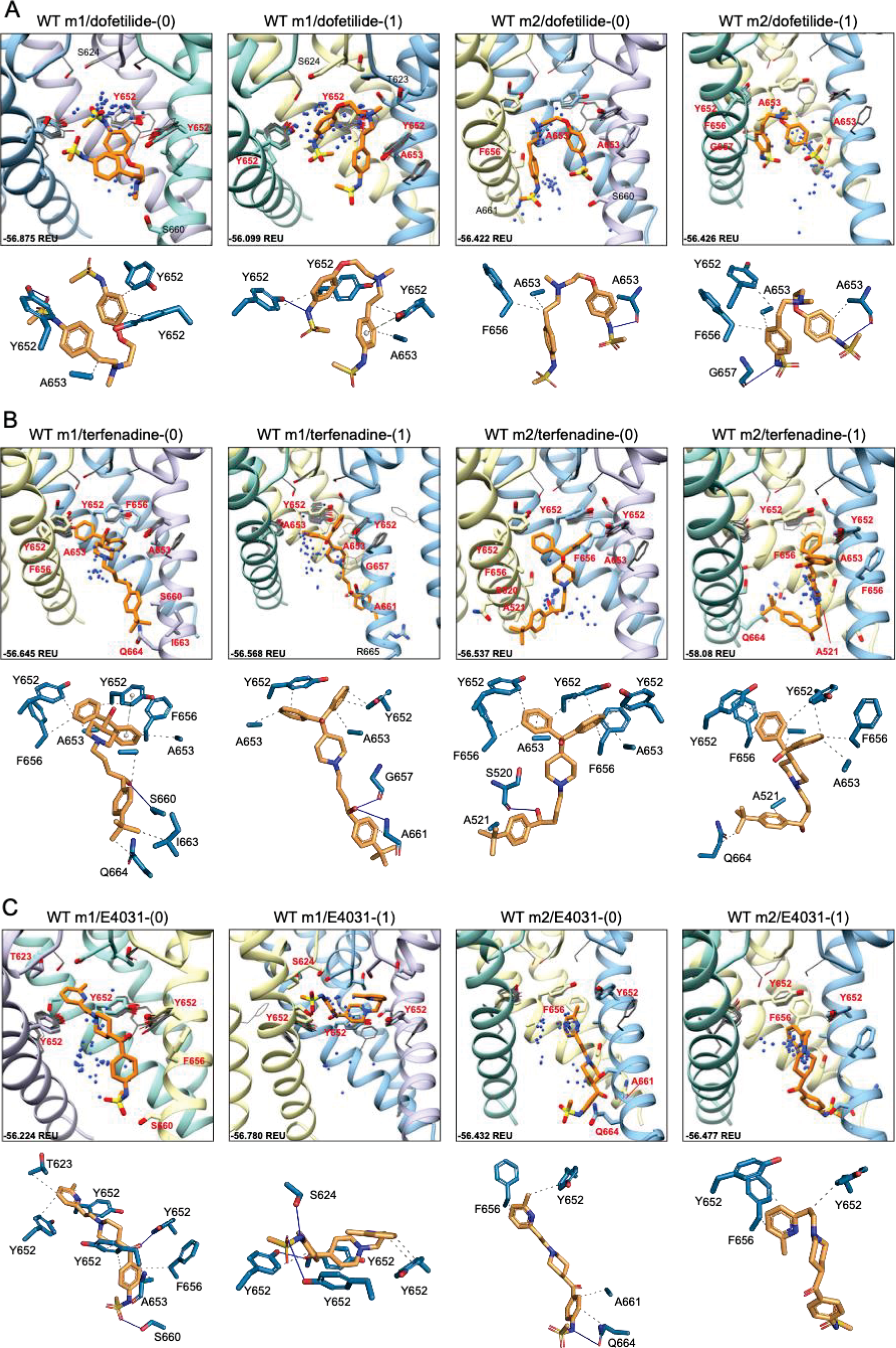
hERG WT drug docking results for (A) dofetilide, (B) terfenadine and (C) E4031. Boxes represent top 50 scoring structures, with top-scoring structure as orange sticks (for carbon atoms) and remaining structures as blue centroids representing central nitrogen/charge group of the respective compound. Oxygen atoms are colored red, nitrogen – blue, sulfur – yellow. Hydrogens are not shown for clarity. Residue contacts to top-scoring structures are shown as sticks and labeled in bold red; residue contacts for all other ligands are shown as gray wires. Chains are colored as in Figure 2.1. Rosetta ligand interface score for each top-scoring structure is shown in Rosetta energy units (REU) in bottom left corner of each box. Protein-ligand interaction profiles (PLIP) for top- scoring poses are shown in lower panels (below boxes) and oriented in approximately the same way. Key contact residues are labeled in black and correspond to bold red labels above. Key residues on protein chains that have been removed for clarity may not appear in upper panels (boxes) but are shown in lower panels for PLIP analysis.

A similar vertical downward shift in the m2 models was observed for the bulkier terfenadine ligand, which was also bound in a more elongated fashion than dofetilide (**Figure 2B**). Most hydrophobic interactions between residues Y652 and F656 involved terfenadine’s hydroxy(diphenyl) functional group, while hydrogen bonding occurred primarily between the hydroxyl group near the tert-butyl tail of terfenadine and backbone amide groups of A661, G657 and S660, the latter also involving its sidechain oxygen in the hERG m1 model with terf-0. Terfenadine also showed a slightly more clustered distribution of the fifty top-scoring models than dofetilide, but with similar positioning of dft-0 closer to the SF in m1 models, and further down in the hERG m2 models.

Lastly, E4031 showed the highest variability in docking modes of all three drugs, although positioning relative to the Y652 plane remained unchanged between m1 and m2 models (**Figure 2C**). Neutral E4031 docked to hERG WT m1 showed a large distribution among the top fifty models, with the top model showing an elongated conformation and protruding its methyl-pyridyl group to T623 near the bottom of the P-helix. On the other hand, charged E4031 models were loosely clustered between the SF and the Y652 plane, with the top-scoring ligand hydrogen-bonding to S624 and Y652. In the hERG WT m2 models, the overall shift of docked ligands went below the Y652 plane and, like in hERG WT m1, neutral E4031 showed a more varied distribution across the hERG pore, whereas charged E4031 was more clustered. Interestingly, the only interactions of the top-scoring charged E4031 - hERG m2 model involved several hydrophobic interactions with F656 and Y652, suggesting a more solvent exposed ligand.

### hERG S641A

Because the S641A mutation in hERG is one of only a handful of known mutations to accelerate the onset of inactivation (Bian et al., 2004), we use it to explore conformational changes that may directly or allosterically impede ion conduction and alter drug affinity during inactivation. S641 is positioned near the top of the S6 segment and is nestled in a relatively hydrophobic environment between F617 and S621 on the pore helix (P-helix), P632 on the S5-SF linker within the same chain, as well as Y616 on the P-helix and F627 on the SF of the adjacent chain (**Figure 3A**). S641A m1 models displayed two major conformational differences compared to their m2 counterparts: a lateral shift of F627 into the conduction path axis and the formation of four lateral fenestrations in the pore. In the S641A m1 radial models there is a large-scale lateral movement of F627 into the axis of the conduction pathway, with the F627 sidechain shifted by nearly a full span of the phenylalanine aromatic ring compared to the WT m1 starting structure (**Figure 3B**). This shift of 0.6 – 2.6 Å is associated with decreasing interaction energy between residues A641 and F627 as F627 moves towards the conduction axis (**Supplemental Fig. 2A**). Movement of F627 into the conduction path also correlates well with a vertical downward movement of Y616 on the adjacent segment by up to 1.5 Å, resulting in increased hydrogen bonding (in some cases) to N629 through reorientation of the tyrosine hydroxyl group (**Supplemental Fig. 2A**). Lastly, the movements of F627 and Y616 appear to be facilitated by the absence of sidechain hydrogen bonding between S620 (located on the P-helix just behind the SF) and the SF in all top-scoring m1 radial models, allowing for increased flexibility of the SF backbone accommodating the residue shifts (**Supplemental Fig. 2A**). A similar range of motion can be seen in the hERG m1 regional models, (those using a movemap encompassing the entire pore domain during the RosettaRelax protocol; see Methods) where the top-scoring model shows F627 overlapping that of the hERG WT m1 starting structure (**Figure 3B, m1**). However, while S620 in all top-scoring hERG m1 radial models is pointed away from the SF, S620 is oriented towards it in about half of the top-scoring hERG m1 regional models, where it interacts with the SF backbone at F627 through strong hydrogen bonding of about -2 kcal/mol; as can be expected, this immobilizes the SF backbone and positions F627 and Y616 closer to their WT starting structure orientations (**Figure 3B, m1 regional SF**; (**Supplemental Fig. 2A**). This effect is markedly more prominent across all m2 models, where S620 hydrogen bonds with the F627 backbone, resulting in a maximal shift of F627 of 0.5 Å (**Figure 3B, m2**; **Supplemental Fig. 2B**).

**Fig. 3.**
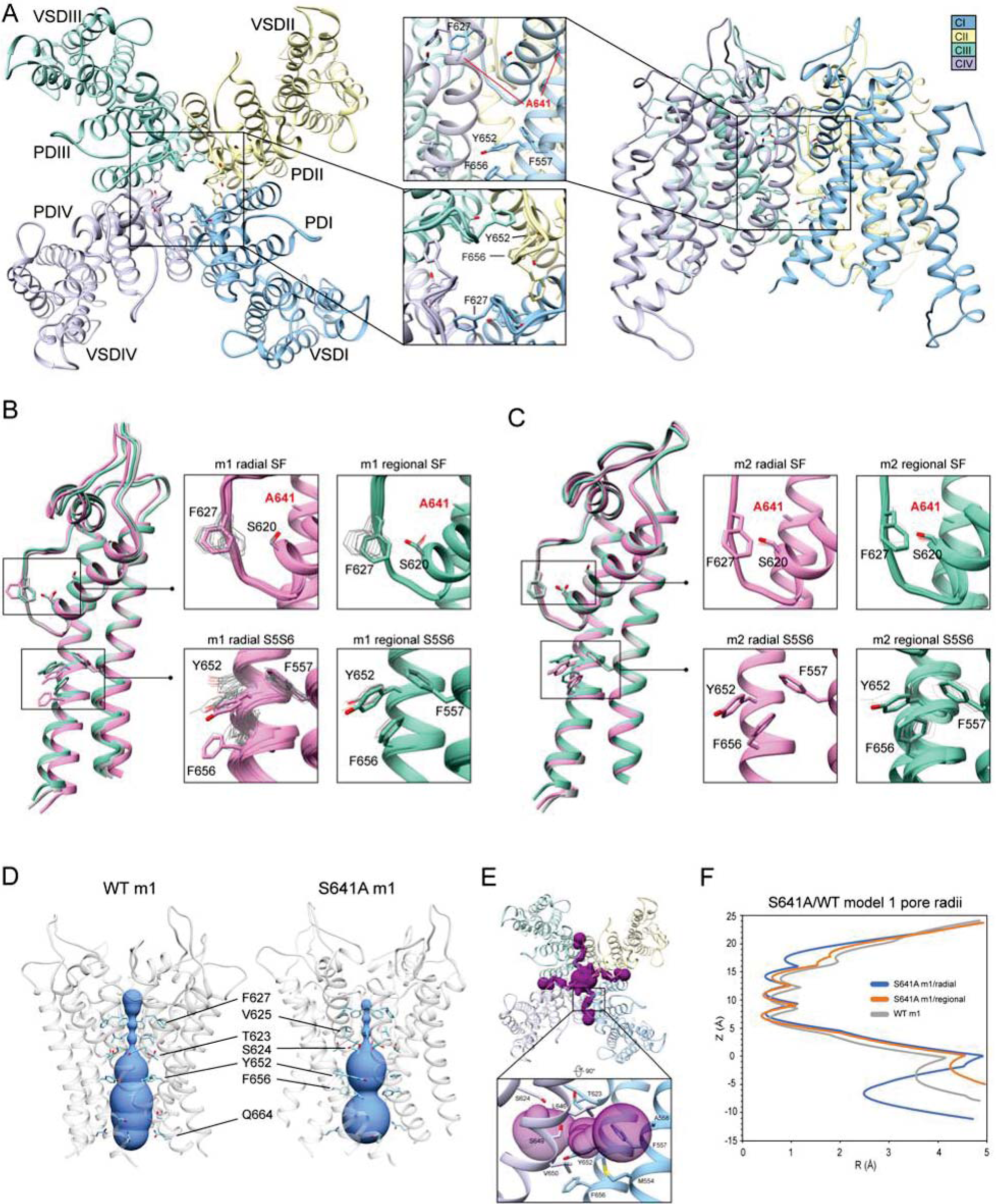
hERG S641A model 1 (m1) and model 2 (m2) comparison. (A) Domain structure of hERG S641A. Chains are colored according to legend. PD: pore domain, VSD: voltage sensing domain. (B) Overlay of hERG S641A m1 models using the radial (rad, pink) and regional (reg, green) movemaps. WT hERG model structure is in gray. Single domain overlay on the left shows the 8^th^ top-scoring structure for S641A m1 (top-scoring for all other models), whereas inset details on the right show top 50 representative structures. Top structure residues in inset detail are shown as sticks, with remaining shown as gray wires. Mutant residues are labeled in bold red. (C) Overlay of hERG S641A m2 models using the radial (rad, pink) and regional (reg, green) movemaps. Depictions are identical to C. (D) hERG WT m1 and hERG S641A m1 model pore volume profile comparison. Volume spheres are in blue, with surrounding residues in light blue. (E) hERG S641A m1 fenestration model top-view, with detail inset rotated by 90 degrees. Fenestration tunnel (purple spheres) are generated by MOLE (Pravda et al., 2018). Fenestration-lining residues are labeled and shown in stick, with chains colored as before. (F) Pore radii profiles are generated by the HOLE program for hERG S641A m1 radial (blue) and regional (orange), WT m1 (gray) models.

The degree of conformational variability can also be seen in the hydrophobic bundle formed by F557, Y652, and F656 on the S5 and S6 segments (**Figure 3B, C, S5S6**). In the hERG m1 models, Y652 displays large vertical movements, parallel to the conduction axis, while F656 undergoes large-scale rotation of the phenyl group and swings laterally into the pore, away from the S5 segment. It is the large-scale movements in this region that form fenestrations in the S641A mutant (**Figure 3E**), showing a clear disengagement of F656 from this hydrophobic cluster in two of the top twenty models (**Figure 3B, m1 radial detail**). Here, interaction energies between F656 and F557 completely absent, with a concomitant disruption of hydrogen bonding between Y652 and S649 on the adjacent chain in one of the S641A mutant models. Interestingly, however, the S649-Y652 hydrogen bond is absent in nearly all S641A models (and about -1.3 to -1.7 kcal/mol when present) despite increased stabilization between F557 and F656 in m1 regional and m2 radial models; it is unclear why the hERG m2 regional models display larger conformational movements (**Figure 3B, C, S5S6; Supplemental Fig. 2B**). In comparison, all WT m1 and m2 relaxed models show strong S649-Y652 hydrogen bonding of about -2.7 kcal/mol, with stabilized interactions between F557 and F656 across all but WT m1 radial models, where an upward shift of S649 and Y652 allows F656 to move outwards towards the channel pore, disrupting its interaction with F557 and facilitating conformational freedom (**Supplemental Fig. 2A,B**). This would suggest that increased movement of the S5 and S6 segments is required to initiate conformational changes in F656, which would then disrupt interactions with Y652 and F557. Thus, this disengagement of residues inside the hydrophobic cluster would enable an outward movement of S5 and S6, expanding the volume of the pore cavity, widening the interaction distance between Y652, F656 and F557, ultimately enabling the formation of the fenestrations (**Figure 3D, F**). Although the chosen S641A model shows a large conformational shift for F656, favorable Lennard-Jones (LJ) and Ramachandran scores give the residue an overall favorable total Rosetta score of -1.21 REU.

Because hERG lacks the stabilizing hydrogen bond network at the SF that is present in other K^+^ channels, such as E71 and D80 in KcsA (Vandenberg et al., 2012), residues at the equivalent positions in hERG (S620 and N629, respectively) affect SF stability in alternate ways. From a mechanistic viewpoint, inactivation through collapse of the SF in our S641A mutant models is primarily facilitated by loss of hydrogen bonding between the S620 hydroxyl oxygen and the SF backbone at F627, with a concomitant shift of F627 into the conduction path, a rearrangement of Y616, and a general increase in hydrogen bonding between Y616 and N629; Y616 being a hydrogen bond partner to N629 in all models including WT. While we did not observe the spontaneous formation of a direct S620-N629 hydrogen bond as in previous MD studies (Kopfer et al., 2012; Schmidtke et al., 2014), we utilized a 1.8 ± 2 Å harmonic constraint between the gamma hydrogen (HG1) and delta oxygen (OD1) of S620 and N629, respectively, to induce hydrogen bonding between the two residues during RosettaRelax in WT m1 rad (**Supplemental Fig. 2C**) and S641A m1 reg (**Supplemental Fig. 2D**) structures. In the WT model, this resulted in hydrogen bonding between Y616 and S641, and a reduction of approximately 4° and 40° in the C-alpha to C-gamma angle and C-alpha to C-delta1 torsion angles of Y616, respectively; these angles were reduced by approximately 6° and 50°, respectively, in the S641A mutant. Although the shift of F627 into the conduction pathway was approximately identical (2.8 Å for WT, 3.0 Å for S641A) in both sets of models, the larger reduction in angles in the S641A mutant would suggest a stabilizing effect of S641 in this potential hydrogen bond network. Using static models, however, it is unclear whether the hydrogen bonds would normally be direct or water mediated.

### Drug docking to hERG S641A

In an attempt to elucidate potential differences in drug docking to the inactivated state, all drugs were docked to the S641A model with fenestrations and the top-scoring S641A model for the hERG m1 and m2 versions, respectively. As can be seen in **Figure 4A**, dofetilide docking deviates significantly from WT in the hERG m1 and m2 models, with top-scoring neutral dofetilide models entering the fenestration region with their sulfonamide moieties. Here, Y652 shifts into the pore cavity to open a space for dofetilide, and F656 shifts back into the hydrophobic region to stabilize the ligand through π-π interactions. Further ligand stabilization through hydrophobic interactions comes from F557 on S5, T623 near the bottom of the SF, and Y652 and A653 on S6 of the adjacent chain. This ligand interaction within the fenestration-forming residues is identical for five out of the 50 top-scoring neutral dofetilide models. For charged dofetilide, only 1 out of 50 models entered the fenestration. It is also interesting to note the high level of conformational variability in Y652 and F656 in the hERG m1 models, a direct result of disengagement of those residues from each other and their interaction with other residues such as F557 in that cluster. Unlike in the hERG m1 models, however, dft-0 and dft-1 docked to S641A hERG m2 models bound similarly to WT.

**Fig. 4.**
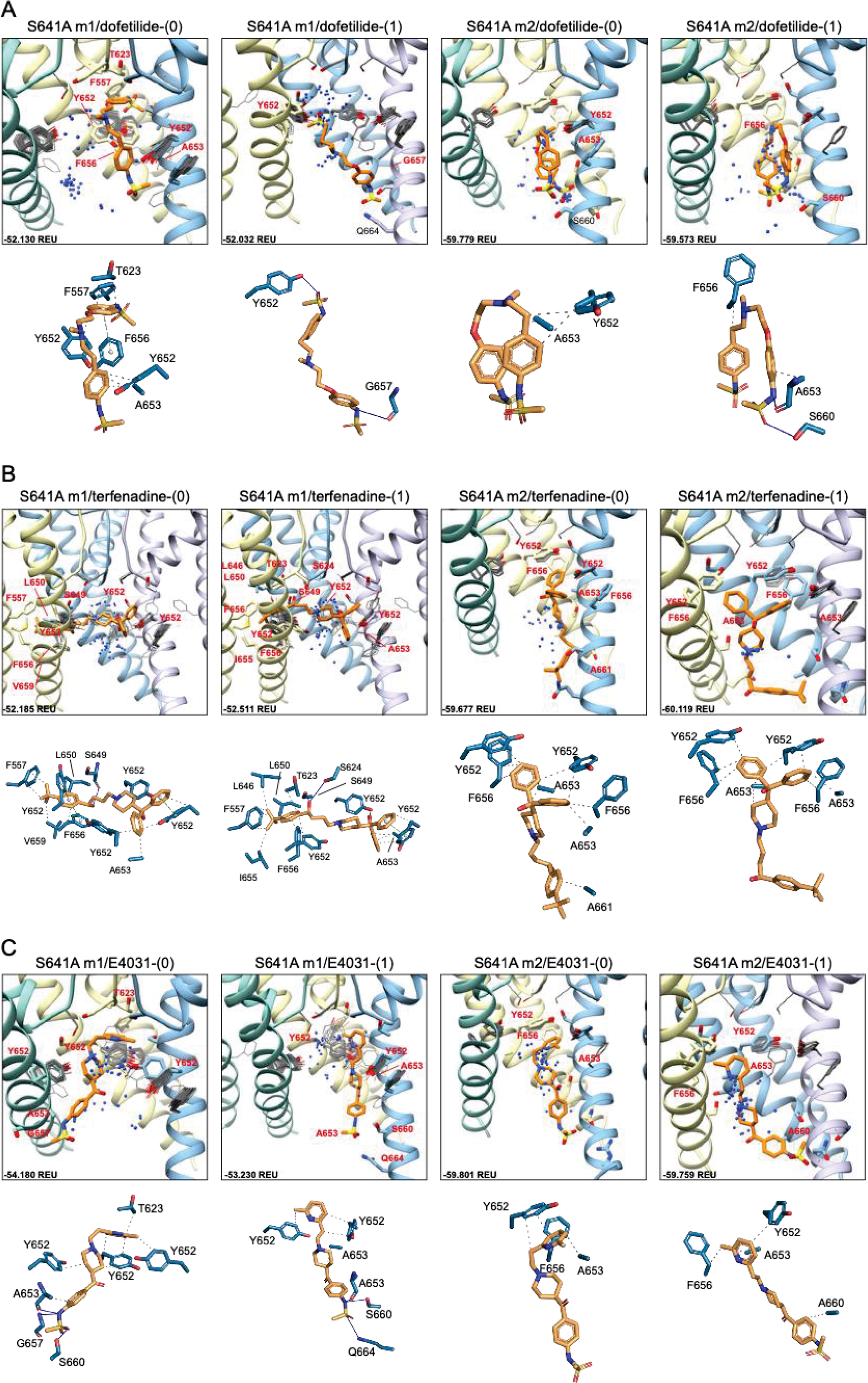
hERG S641A drug docking results for (A) dofetilide, (B) terfenadine and (C) E4031. Boxes represent top 50 scoring structures, with top-scoring structure as orange sticks (for carbon atoms) and remaining structures as blue centroids representing central nitrogen/charge group of the respective compound. Oxygen atoms are colored red, nitrogen – blue, sulfur – yellow. Hydrogens are not shown for clarity. Residue contacts to top-scoring structures are shown as sticks and labeled in bold red; residue contacts for all other ligands are shown as gray wires. Chains are colored as in Figure 2.1. Rosetta ligand interface score for each top-scoring structure is shown in Rosetta energy units (REU) in bottom left corner of each box. Protein-ligand interaction profilers (PLIP) for top- scoring poses are shown in lower panels (below boxes) and oriented in approximately the same way. Key contact residues are labeled in black and correspond to bold red labels above. Key residues on protein chains that have been removed for clarity may not appear in upper panels (boxes) but are shown in lower panels for PLIP analysis.

For neutral and charged terfenadine docked to the S641A hERG m1 model, 15 and 16 out of 50 top-scoring terf-0 and terf-1 models, respectively, entered the fenestration region (**Figure 4B**). All terfenadine ligands entering the fenestration do so through their *tert*-butyl functional groups, with the directly attached benzene ring stabilized either by π-π interactions with Y652 and/or F656, or general hydrophobic interactions, and through hydrogen bonding of the hydroxyl directly after the *tert-*butyl group with S624 on the SF, or S649 on S6 of the adjacent domain. Unlike dofetilide, terfenadine can span the full diameter of the pore cavity, enabling additional π-π, hydrophobic, and hydrogen bonding interactions with residues of the domain directly across the pore. Additionally, because terfenadine does not have the charged sulfonamide functional group, it is capable of binding deeper inside the fenestration and interacting with more residues lining it. Similar to dofetilide, large conformational changes can be seen in key residues Y652 and F656 indicating highly variable interactions in the S641A hERG m1 models with fenestrations, whereas ligand-protein interactions in the hERG m2 models are similar to WT.

E4031 exhibited a slightly different mode of binding within the fenestration. Although none of the top models entered the fenestration (**Figure 4C**), 16 and 18 out of the top 50 neutral and charged models, respectively, exhibited minimal interaction with F557. Interestingly however, only 2 neutral E4031 models, and 5 charged E4031 models, penetrated the fenestration deep enough for strong interactions with F557. Primary interactions involved π-π or hydrophobic interactions of the pyridyl functional group with the top portion of Y652, with the sulfonamide end stabilized through hydrogen bonding and hydrophobic interactions with a variety of residues on adjacent and opposite domains. In some cases (primarily with charged E4031), Y652 shifted either laterally into the pore cavity or vertically towards the SF allowing the pyridyl group to penetrate all the way to F557 where it was able to form strong π-π and hydrophobic interactions with F557 and F656 and hydrophobic residues lining the region. Additionally, in a single instance for neutral E4031, the sulfonamide group was able to penetrate the fenestration making contact with F557. Similar to dofetilide and terfenadine, binding of E4031 to S641A m1 models showed conformationally dynamic Y652 and F656 residues compared to m2 models. Additionally, both neutral and charged E4031 models had a much higher propensity for assuming a more linear conformation in S641A hERG m2 models, binding below the Y652 plane and aligning themselves more parallel to the S6 segments and occluding the intracellular gate.

Overall, the fenestration region observed in our S641A models provides an alternative binding site for aliphatic/partially aliphatic and aromatic moieties of drugs entering the pore cavity and correlates with a proposed hERG inactivation. Notably, while we demonstrate that this region is involved in binding of all drugs, there is a larger number of top models with terfenadine compared to dofetilide entering the fenestration region, with a preference of the pyridyl versus sulfonamide group; observations that are both in agreement with and contrary to existing experimental evidence (Ficker et al., 2001; Herzberg et al., 1998; Mitcheson and Perry, 2003; Perrin et al., 2008; Thouta et al., 2018; Wu et al., 2015).

### hERG S641T

While smaller aliphatic substitutions of S641 facilitate C-type inactivation, bulkier charged and polar groups have been shown to disrupt it (Bian et al., 2004). To explore a potential mechanism for this phenomenon, we use the S641T mutation, which replaces a serine with a bulkier threonine, maintaining the aliphatic nature of the S641A mutation, in conjunction with the hydroxyl group of the WT serine (**Figure 5A**). Conformational variability of key residues F627, Y652 and F656 in all S641T models was consistent with usage of WT m1 (larger-scale movements) and m2 (smaller-scale movements) starting structures, though no opening of the hydrophobic pocket was observed (**Figure 5B, C**). However, a number of interesting differences arose in the SF region when comparing the two sets of models. For S641T hERG m1, the orientation of T641 remains identical in both radial and regional models, with the methyl moiety of threonine pointed downwards towards F627, causing the latter to shift into the conduction path (**Figure 5B, D, m1 models**). The extent of this shift, however, is dependent on the orientation of the S620 hydroxyl oxygen. In the case where the oxygen is pointing towards the SF, it forms hydrogen bonds with the F627 NH group, stabilizing the backbone and shifting F627 about 1.3 Å into the conduction path. Where S620 points away from the SF, that distance is nearly doubled (**Supplemental Fig. 3A, top**). As a result, interaction energies between T641 and F627 are around -0.9 kcal/mol at the larger distances, and about -1.2 kcal/mol when the F627 aromatic ring is closer to T641. The stronger (more negative) interaction energy is due to favorable desolvation and attraction terms (*fa_sol* and *fa_atr*, respectively) characteristic of a hydrophobic environment, albeit with much higher repulsive energy (*fa_rep*) due to steric clash with the T641 methyl group. This is maintained in m1 regional models, where S620 points away from the SF, thereby allowing movement of F627 into the conduction path by 1.7 - 2.9 Å, with the same pattern of interaction energy values, and ultimately leading to narrowing of the SF (**Figure 5C, D, E; Supplemental Fig. 3A, bottom**).

**Fig. 5.**
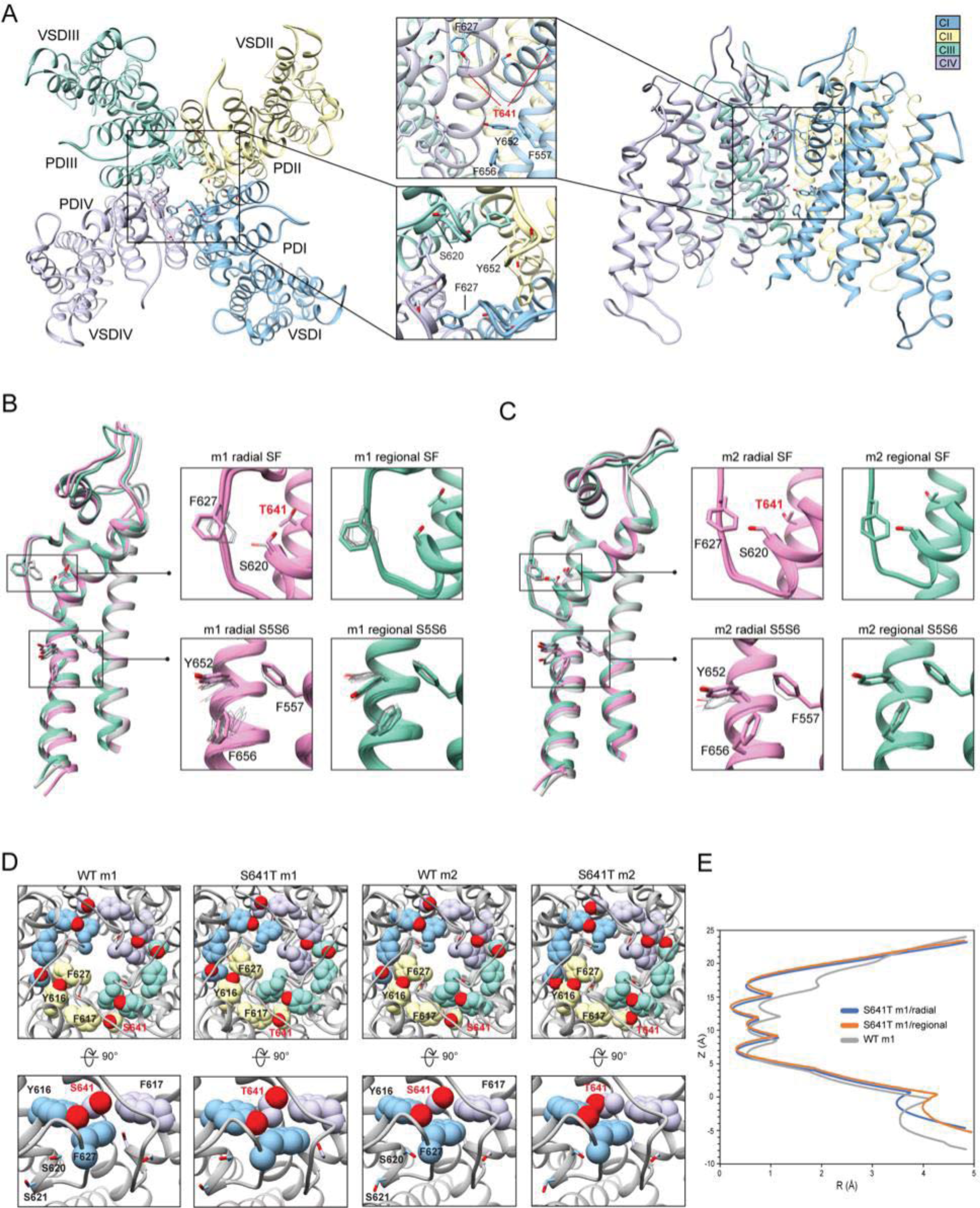
hERG S641T model 1 (m1) and model 2 (m2) comparison. (A) Domain structure of hERG S641T mutant models. Chains are colored according to the legend. PD: pore domain, VSD: voltage sensing domain. (B) Overlay of hERG S641T m1 models using the radial (rad, pink) and regional (reg, green) movemaps. WT model structure is in gray. Single domain overlay shows top-scoring structures, whereas inset details show top 50 structures. Top structure in inset detail is shown as stick, with remaining structures shown as gray wires. Mutant residue Is labeled in bold red. (C) Overlay of hERG S641T m2 models using the radial (rad, pink) and regional (reg, green) movemaps. Depictions are identical to B. (D) Comparison of residue 641 (labeled bold red) and associated residue positions surrounding the selectivity filter (SF) for hERG WT and S641T m1/m2 models. Top row: top view of hERG channel pore, bottom row: side-view of S/T641 interaction details. Residues are shown as spheres and colored as in Figure 2.1. (E) Pore radii generated by the HOLE program for hERG S641T m1 radial (blue) and regional (orange) as well as WT m1 (gray) models.

In the case of S641T hERG m2 models, the T641 residue is rotated about 90° counterclockwise, with the methyl group pointing towards F617, leaving the region open for F627, which retains an orientation similar to WT, with a lateral shift of only about 0.5 Å across all m2 models (**Figure 5D, C; Supplemental Fig. 3B**). While this is in large part due to strong hydrogen bonding between S620 and the SF backbone, the orientation of T641 provides a similar level of favorable desolvation energy without the large steric clash with the methyl group as in the m1 models, giving a stronger interaction between T641 and F627 with an average of -1.46 kcal/mol in m2 models versus -0.88 kcal/mol in m1 models (**Supplemental Fig. 3B**). Additionally, while the interaction energy between T641 and the nearby F617 residue remain similar between all models (with a slight penalty for steric repulsion in m2 models), interaction between T641 and Y616 on the neighboring chain vary greatly. In m1 models, this interaction is on average about -0.7 kcal/mol, stemming from a combination of attractive and repulsive forces between aliphatic and polar moieties; in m2 models, there appears to be virtually no interaction between the two residues (**Supplemental Fig. 3B**). The overall effect of this is that both Y616 and F617 are drawn closer to T641, crowding out F627 in the m1 models; the reverse is true in the m2 models, which provide a hydrophobic pocket for F627. From the perspective of attenuating inactivation, the most probable threonine orientation in the S641T mutation is the one found in the hERG m2 models, where the positions of F617 and Y616 on the adjacent chains enable F627 to remain locked in position within this hydrophobic pocket.

### Drug docking to hERG S641T

Docking of dofetilide, terfenadine and E4031 to hERG S641T showed similar protein-ligand interactions to the WT models (**Figure 6**). For dofetilide, top-scoring models retained their bent structure in both hERG m1 and m2 models, similar to WT, with a large proportion of hERG m1 models reaching above the Y652 plane with associated hydrophobic, π-π, and hydrogen bonding interactions with Y652 and SF residues (**Figure 6A**). The reverse is true for hERG m2 models, with the majority of ligands, including dft- 0 and dft-1 top-scoring models, binding near the intracellular gate. Similar observations can be made for terfenadine, with the notable exception that both neutral and charged terfenadine ligands in the m1 models bound Y652 residues with their *tert*-butyl groups, as opposed to the branched diphenyl end seen in the hERG m2 and WT models, enabling a deeper penetration of the region below the SF for the top-scoring neutral terfenadine model (**Figure 6B**). Lastly, for E4031 models, both the overall ligand distribution within the hERG cavity and protein-ligand binding interactions of the top-scoring models were nearly identical to WT (**Figure 6C**). The similarity of drug binding to WT suggests that the S641T mutation preserves overall residue orientation within the pore cavity, while rearranging residues at the SF to maintain ion conduction.

**Fig. 6.**
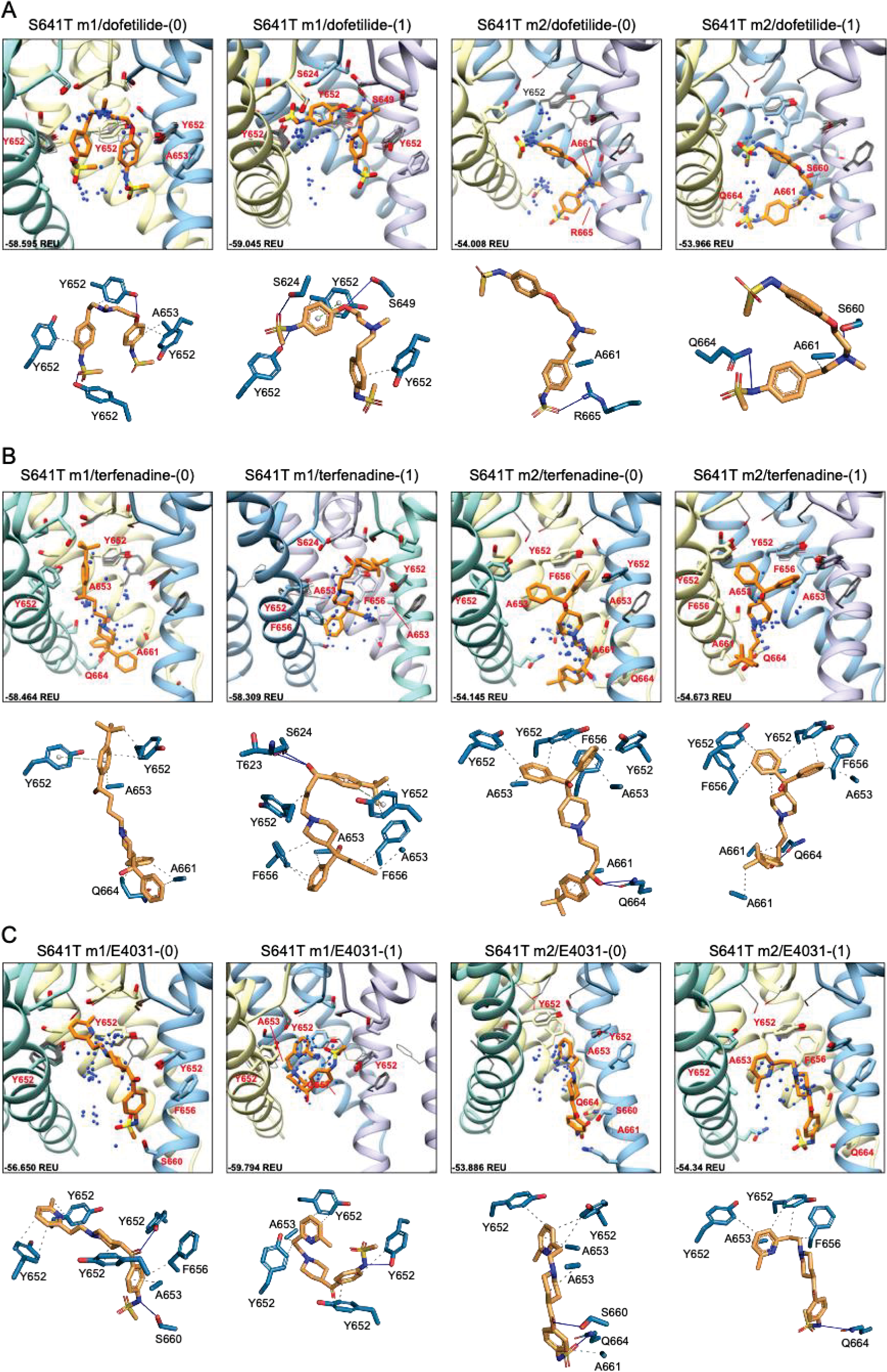
hERG S641T drug docking results for (A) dofetilide, (B) terfenadine and (C) E4031. Boxes represent top 50 scoring structures, with top-scoring structure as orange sticks (for carbon atoms) and remaining structures as blue centroids representing central nitrogen/charge group of the respective compound. Oxygen atoms are colored red, nitrogen – blue, sulfur – yellow. Hydrogens are not shown for clarity. Residue contacts to top-scoring structures are shown as sticks and labeled in bold red; residue contacts for all other ligands are shown as gray wires. Chains are colored as in Figure 2.1. Rosetta ligand interface score for each top-scoring structure is shown in Rosetta energy units (REU) in bottom left corner of each box. Protein-ligand interaction profilers (PLIP) for top- scoring poses are shown in lower panels (below boxes) and oriented in approximately the same way. Key contact residues are labeled in black and correspond to bold red labels above. Key residues on protein chains that have been removed for clarity may not appear in upper panels (boxes) but are shown in lower panels for PLIP analysis.

### hERG S620T

Because the S641T mutation is distal to the SF, we also tested the hERG S620T mutation, which has been shown to strongly attenuate inactivation (Herzberg et al., 1998), and replaces a serine with threonine at position 620 on the P helix just behind the SF (**Figure 7A**). Molecular dynamics studies suggest that the mutation introduces steric hindrance behind the SF through threonine’s methyl group, the consequence of which may, firstly, disrupt a potential hydrogen bond between S620 and N629 – critical to the collapse of the SF during inactivation – and, secondly, introduce steric hindrance that prevents F627 from shifting into the conduction axis during inactivation (Kopfer et al., 2012; Li et al., 2021; Stansfeld et al., 2008) (**see Figure 5, S641T**). As we hypothesized, conformational variation based on overall RMSD values, in hERG m2 models was significantly lower than in hERG m1, with all critical residues (F627, Y652, F656 and F557) in the top-scoring models retaining near-identical orientations with respect to each other, suggesting that the m2 models are more conformationally stable (**Figure 7B, C insets**). In our models, S620T remains in a near identical orientation in all top-scoring models, with the hydroxyl oxygen pointed towards the backbone of F627 with strong hydrogen bonding (around -2 kcal/mol for both m1 and m2 models), and the methyl group directed towards F627 of the adjacent chain, stabilized through hydrophobic interactions (**Figure 7B-D; Supplemental Fig. 4**). We do however see a slight shift in F627 into the conduction pathway in both sets of models, potentially due to the bulkiness of the threonine at the position 620 (**Figure 7B, domain overlay, 7E**). In hERG S620T m1 models, this shift ranges from 1 to 1.6 Å, whereas it is only about 0.5 Å in all m2 models (**Figure 7C, domain overlay**, **Supplemental Fig. 4A**). Additionally, a much larger sampling of conformational space can be seen in residues Y652 and F656 on the S6 segment of the hERG S620T m1 models compared to those in the m2 set, analogous to the effects seen in the S641A/T mutations. Compared to WT models, the presence of strong hydrogen bonding between the hydroxyl oxygen at the 620 position and the F627 backbone correlates to that found in WT m2 models, while hydrophobic interaction (measured by desolvation energy *fa_sol*) between T620 and the neighboring F627 residue was closer to WT m2 models, suggesting that the m2 model is more amenable to the hERG open state (**Figure S4B**). Lastly, although no hydrogen bond contacts could be detected between S620 and N629 in the WT models, the bulky methyl group of the T620 mutant clearly stabilized the position of F627 on the neighboring chain in both hERG m1 and m2 models, supporting the idea of a physical obstruction of F627 preventing from shifting towards the conduction axis during inactivation.

**Fig. 7.**
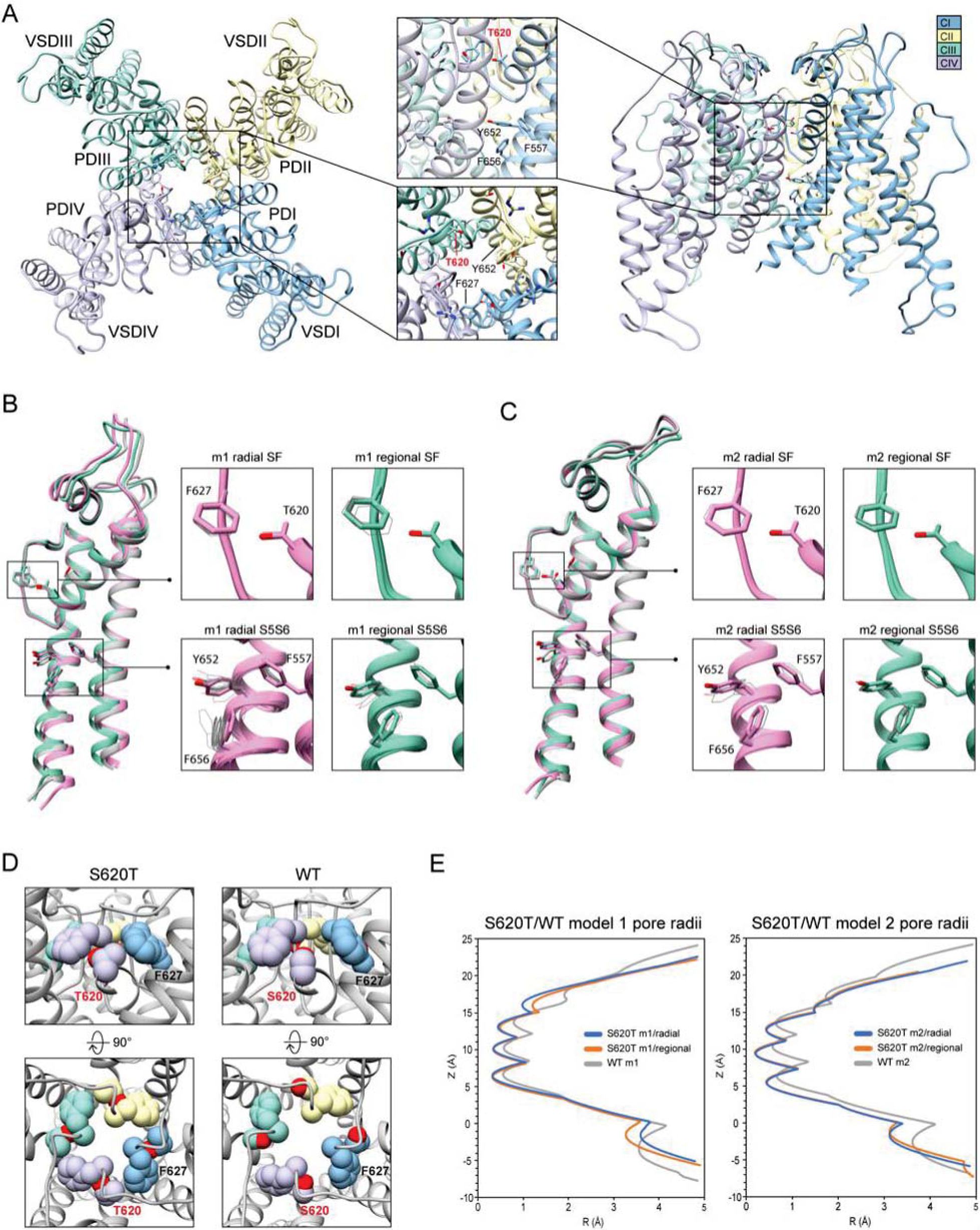
hERG S620T model 1 (m1) and model 2 (m2) comparison. (A) Domain structure of hERG S620T. Chains are colored according to legend. PD: pore domain, VSD: voltage sensing domain. (B) Overlay of hERG S620T m1 models using the radial (rad, pink) and regional (reg, green) movemaps. WT model structure is in gray. Single domain overlay shows top-scoring structures, whereas inset details show top 50 structures. Top structure in inset detail is shown as sticks, with remaining shown as gray wires. Mutant residue is labeled in bold red. (C) Overlay of hERG S620T m2 models using the radial (rad, pink) and regional (reg, green) movemaps. Depictions are identical to B. (D) Comparison of residue 620 (labeled bold red) and associated residue positions surrounding the SF in hERG WT and S620T models. Top row: detailed side-view of residue 620 interactions, bottom row: top view of hERG channel pore. Residues are shown as spheres and colored according to their respective chains (see legend in A). (E) Pore radii, *R*, profiles along the *z* axis are generated by the HOLE program for hERG S620T m1 radial (blue) and regional (orange), WT m1 (gray) models.

### Drug docking to hERG S620T

Dofetilide docking to S620T hERG m1 models was significantly different from WT, as both dft-0 and dft-1 ligands bound in a more elongated manner, with simultaneous sulfonamide hydrogen bond contacts to one or two Q664 residues near the intracellular gate, and hydrogen bonds to S649/S624 near and at the bottom of the SF (**Figure 8A**). For dofetilide, ligand distributions were relatively dispersed, with only minor clustering of dft- 0 near the SF in S620T m1 hERG model, and near the lower end of the pore cavity in S620T hERG m2. Similar to WT, however, hERG m2 models for dofetilide, terfenadine and E4031 all showed the majority of ligands docked below the Y652 plane (**Figure 8A- C, m2 models**).

**Fig. 8.**
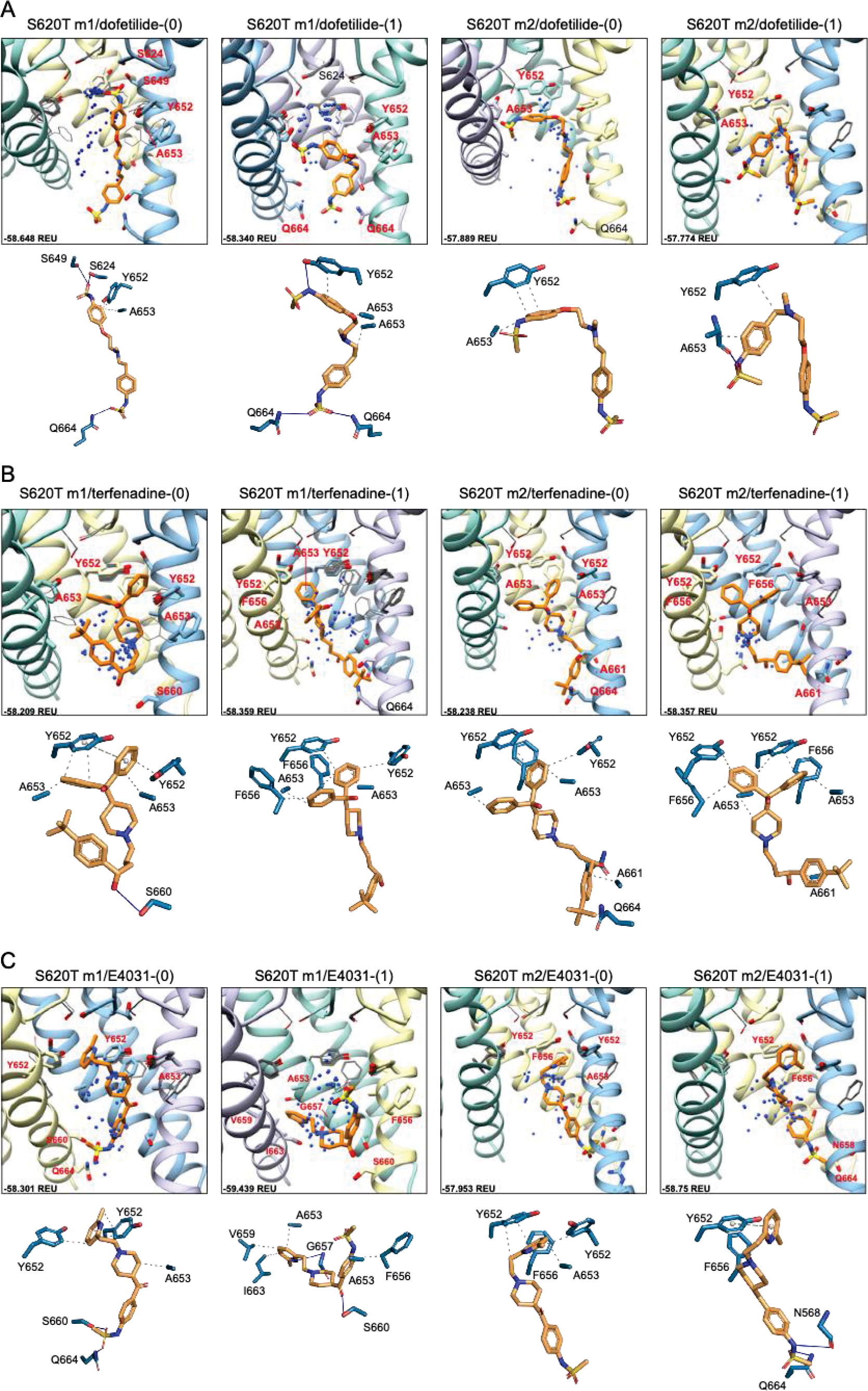
hERG S620T drug docking results for (A) dofetilide, (B) terfenadine and (C) E4031. Boxes represent top 50 scoring structures, with top-scoring structure as orange sticks (for carbon atoms) and remaining structures as blue centroids representing central nitrogen/charge group of the respective compound. Oxygen atoms are colored red, nitrogen – blue, sulfur – yellow. Hydrogens are not shown for clarity. Residue contacts to top-scoring structures are shown as sticks and labeled in bold red; residue contacts for all other ligands are shown as gray wires. Chains are colored as in Figure 2.1. Rosetta ligand interface score for each top-scoring structure is shown in Rosetta energy units (REU) in bottom left corner of each box. Protein-ligand interaction profiles (PLIP) for top- scoring poses are shown in lower panels (below boxes) and oriented in approximately the same way. Key contact residues are labeled in black and correspond to bold red labels above. Key residues on protein chains that have been removed for clarity may not appear in upper panels (boxes) but are shown in lower panels for PLIP analysis.

Similarities could also be seen between WT and S620T hERG docking for terfenadine, with the (diphenyl)hydroxymethyl group exhibiting the bulk of hydrophobic interactions with Y652, F656 and A653, and terfenadine ligands binding lower in the hERG m2 models, as evidenced by the lack of perturbations of Y652 and F656 residues (**Figure 8B, m2 models**). Clustering, however, was more pronounced in the hERG m2 models, with both terf-0 and terf-1 showing a higher propensity for clustered ligands towards the bottom half of the pore cavity. No real clustering could be discerned for E4031 ligands, however, which bound to all models in a very similar way as WT (**Figure 8C**). For all ligands, most interactions were hydrophobic in nature, with only a few of the top-scoring models (terf-0 in S620T m1, and E4031-1 in S620T m2) exhibiting more directional π- π interactions.

### Relative stability of WT and mutant models

To explain structural differences between models 1 and 2, we look at the root mean square deviations (RMSD) between Cα atoms to describe conformational variability. To assess the extent of conformational variability in m1 and m2 models when subjected to a Rosetta FastRelax protocol, RMSD was measured against respective hERG m1/m2 input models for the top 10% (1,000) of generated models using both radial (**Figure 9A**) and regional (**Figure 9B**) movemaps. For both radial and regional models, m1 models showed significantly higher RMSD values than m2 models. However, m1 radial models indicated a higher degree of structural variability than m2 radial models. Additionally, differences in mean RMSD between radial and regional models within the m1 and m2 groups were relatively small for m1 models (RMSD ∼ 0.1 Å), whereas m2 models had nearly double RMSD values using the radial movemap. This adds to the observation that m1 models, with R582 pointed away from the S5P turret, exhibit a higher degree of conformational freedom than their m2 counterparts. Thus, within the context of hERG channel inactivation, structural changes associated with the S641A fast-inactivated mutant with fenestrations may require larger conformational changes within the pore domain.

**Fig. 9.**
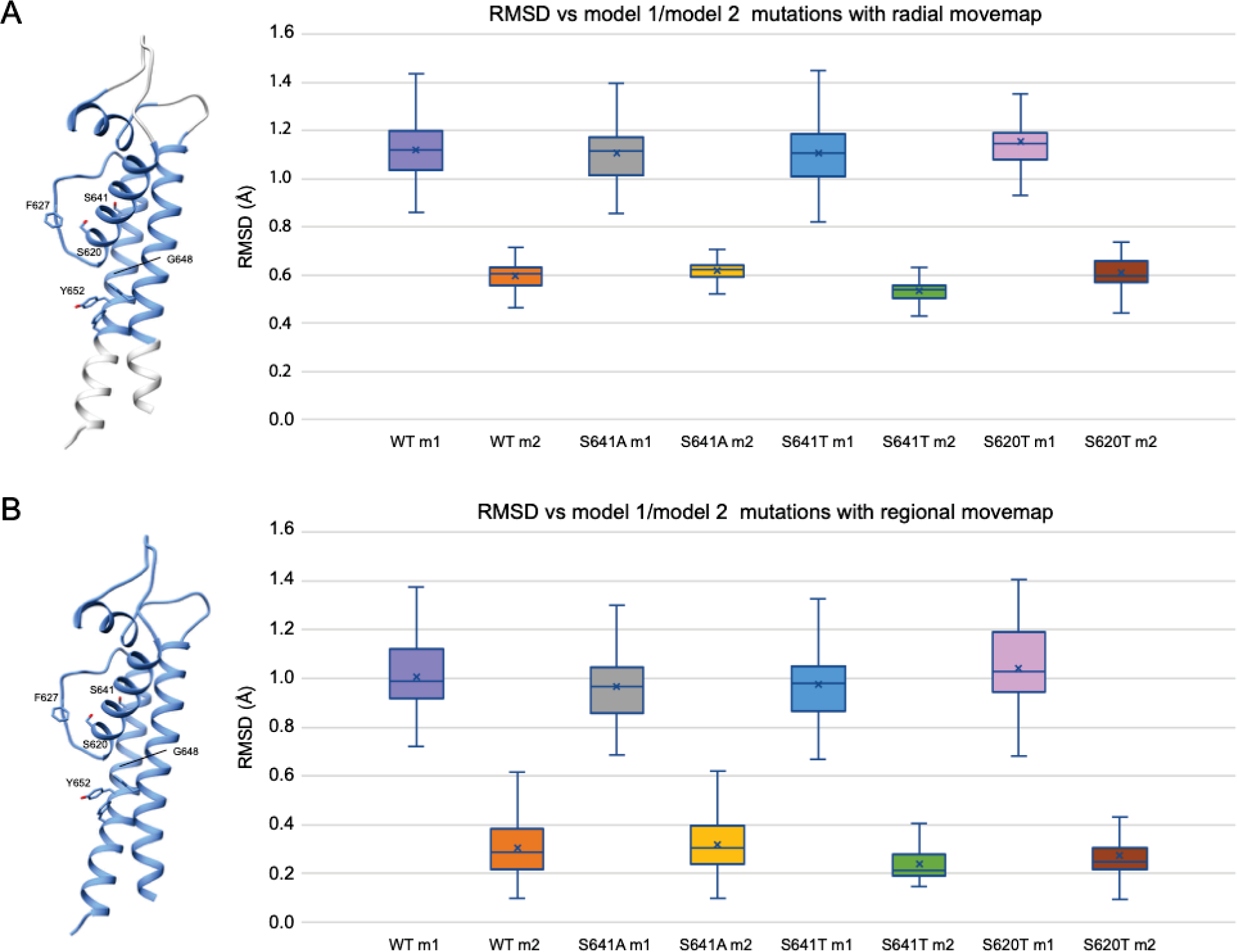
Relative stability of WT and mutant models using the (A) radial and (B) regional movemaps. Single domain representations are shown on left with regions specified as movable by movemap depicted in blue, and key residues labeled. RMSD: Cα root-mean-square deviation in Angstroms. RMSD for m1 and m2 mutant models were measured against m1 and m2 WT hERG channel models, respectively. Error bars represent Max and Min values; Mean (x); Median (horizontal bar).

### Comparison of ligand conformation and interface scores of bound drugs

Although GALigandDock utilizes Rosetta’s latest betanov2016 energy function (Park et al., 2021), which accounts for torsional strain energies of the ligand, in addition to using fitted van der Waals distances, solvation potentials, and partial charges, it was difficult to make conclusions between drug conformation/binding modes and interface scores in our models. Comparative ligand interface scores and probability density plots for all 2000 docked dofetilide, terfenadine, and E4031 structures can be found in **Figures 10**, **11**, and **12**, respectively. On a global level, all drugs bound to WT models had a median interface score of -54 to -55 Rosetta energy units (REU) across m1 and m2 models, with relatively small variability, which can be attributed to a single mode of binding. This was followed by more favorable (more negative) median interface scores within the range of –56 to - 58 REU for all drugs bound to S620T m1/m2 mutants. As would be expected, large differences arose in the S641A models, with m1 models showing significantly less favorable scores (∼ –50 REU) over a smaller range than m2 models where the median scores were approximately –57 REU. For S641T m1 models, median scores were around –56 REU compared to –52 REU for m2 models across all drugs. Overall, WT and S620T models showed the most consistent score distributions across the different mutations between m1 and m2 models, whereas S641A/T had the largest score fluctuations.

**Fig. 10.**
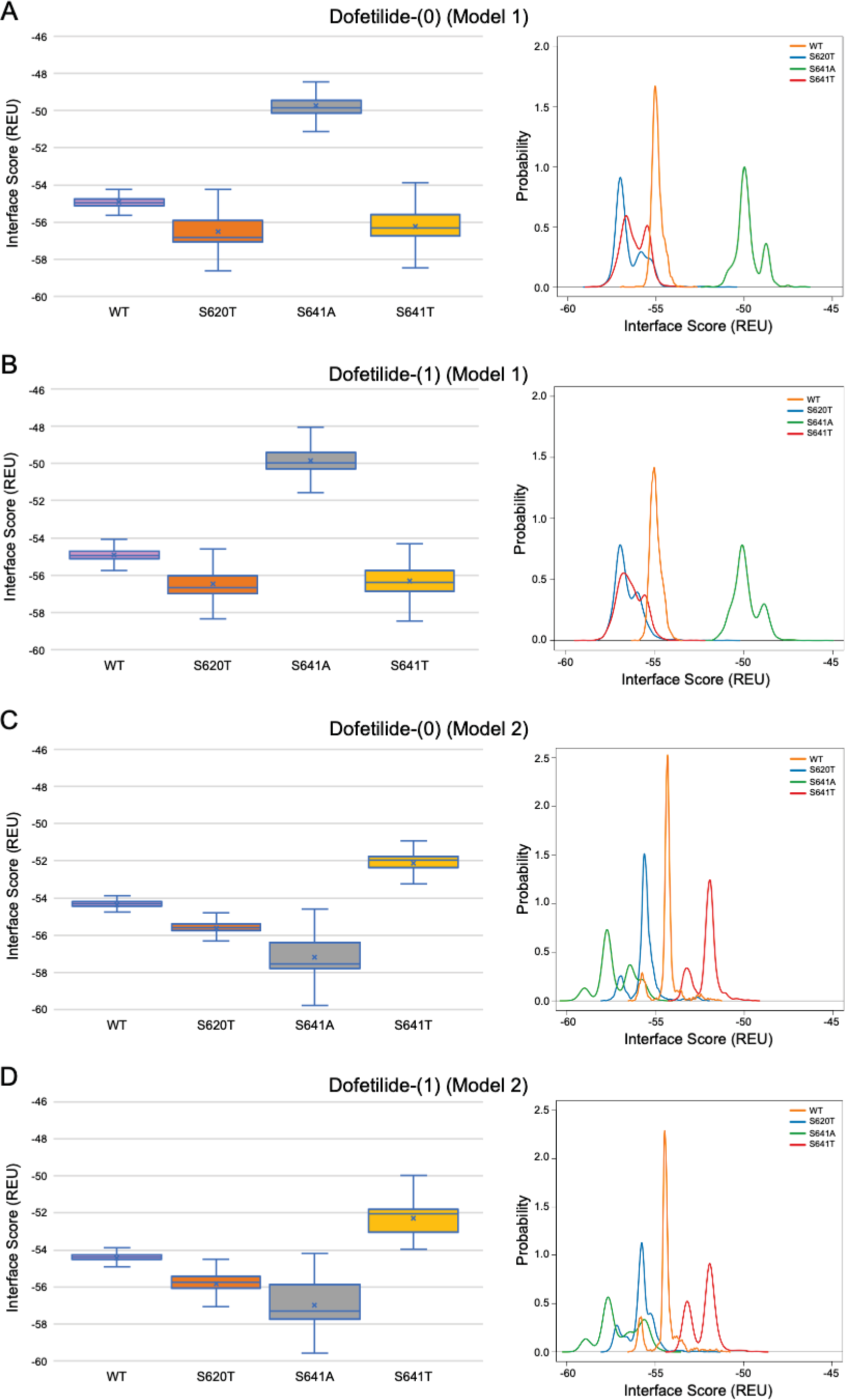
Interface score distributions for hERG channel interactions with dofetilide. Left Panels: Box-and-whisker plots of interface scores in Rosetta energy units (REU). Right Panels: Probability density charts plotting probability density against interface scores (REU) All plots utilize 2,000 generated structures. Plots are defined for (A) neutral (0) model 1, (B) charged (1) model 1, (C) neutral (0) model 2 and (D) charged (1) model 2 structures. Error bars represent Max and Min values; Mean (x); Median (horizontal bar).

**Fig. 11.**
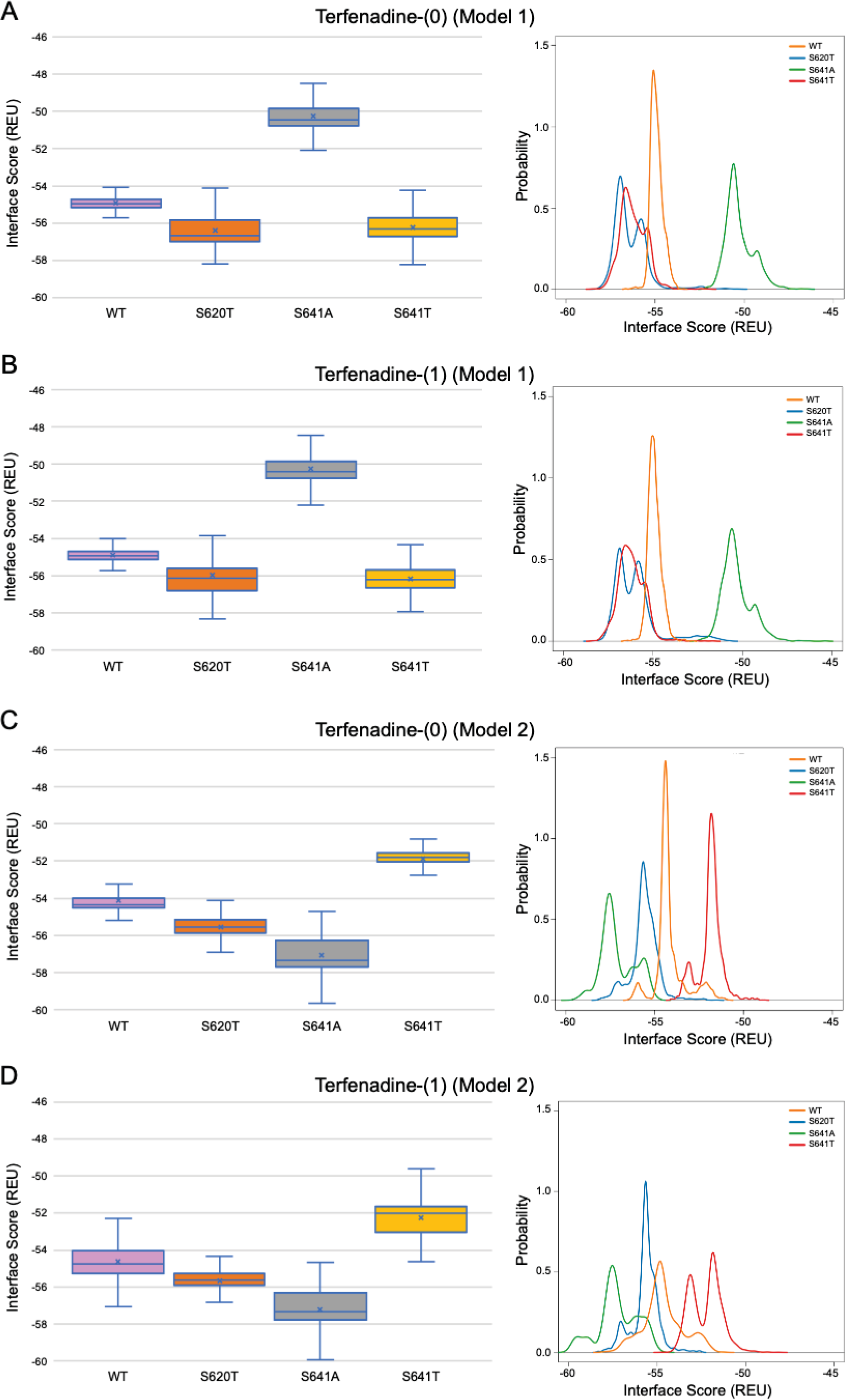
Interface score distributions for hERG channel interactions with terfenadine. Left Panels: Box-and-whisker plots of interface scores in Rosetta energy units (REU). Right Panels: Probability density charts plotting probability density against interface scores (REU). All plots utilize 2,000 generated structures. Plots are defined for (A) neutral (0) model 1, (B) charged (1) model 1, (C) neutral (0) model 2 and (D) charged (1) model 2 structures. Error bars represent Max and Min values; Mean (x); Median (horizontal bar).

**Fig. 12.**
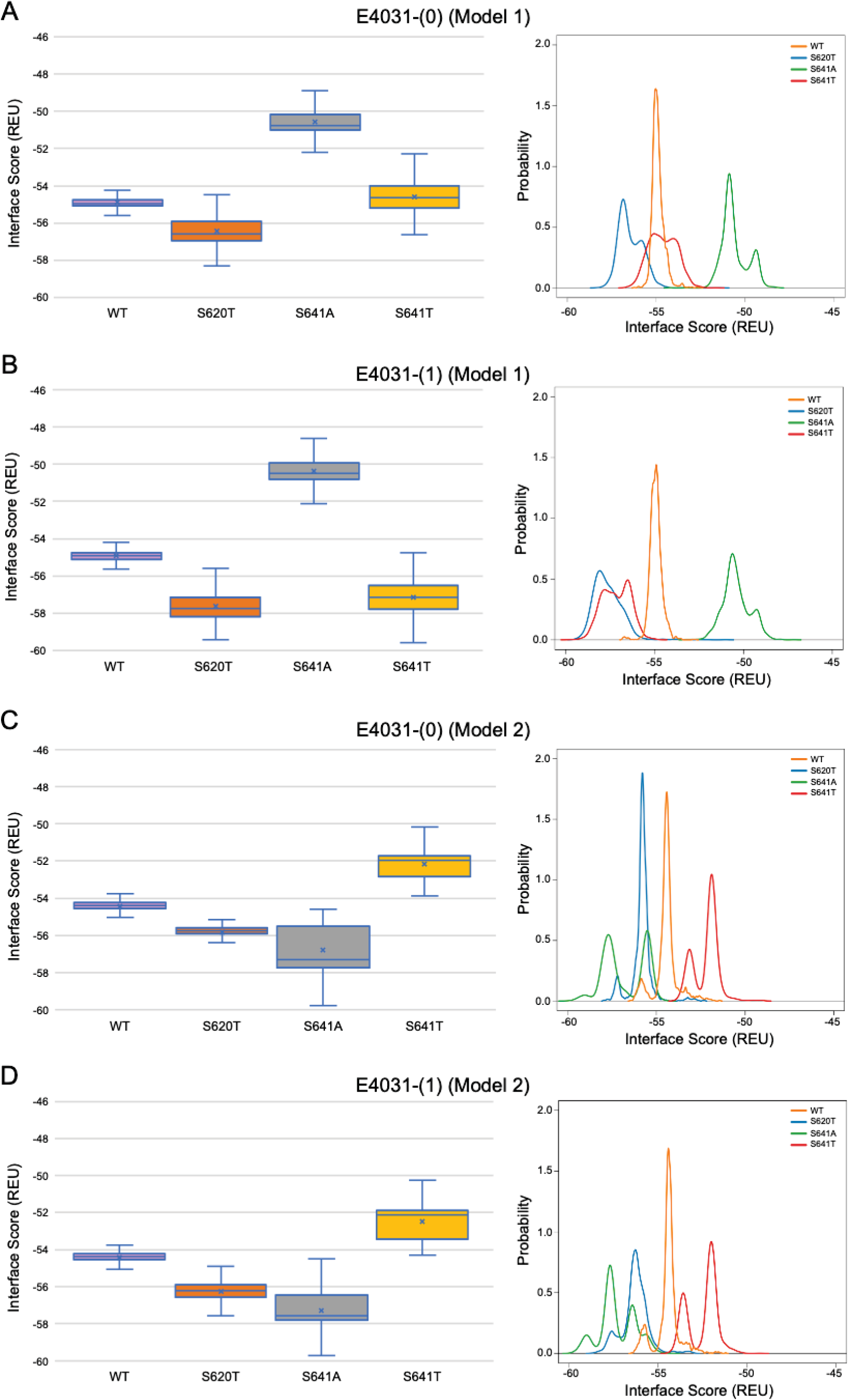
Interface score distributions for hERG channel interactions with dofetilide. Left Panels: Box-and-whisker plots of interface scores in Rosetta energy units (REU). Right Panels: Probability density charts plotting probability density against interface scores (REU). All plots utilize 2,000 generated structures. Plots are defined for (A) neutral (0) model 1, (B) charged (1) model 1, (C) neutral (0) model 2 and (D) charged (1) model 2 structures. Error bars represent Max and Min values; Mean (x); Median (horizontal bar).

While obvious exceptions to this observation exist, such as the more solvent exposed aliphatic moieties of dft-1 in S641T m2 (-53.966 REU) compared to the more buried dft-1 docked to the S641T m1 (-59.045 REU) (**Figure 6A**), the same criteria fall short when comparing terfenadine docked to S641A versus S641T m2 models, both of which have similar levels of protein-ligand interaction and solvent exposure (**Figure 4B and 6B, respectively**). Furthermore, the S641A mutant with fenestrations was the only one that enabled some fraction of all top-scoring drugs (though not necessarily the top pose) to sample an alternate space within the hERG pore (**Figure 4B, terfenadine m1 models)**. Although this alternate docking pose showed an increase in hydrophobic and π-π interactions with a subsequent loss of solvent accessibility and hydrogen bonding, the S641A m1 models had the lowest scores GALigandDock scores of all mutant models, regardless of the type and conformation of docked drug (**Figure 4A-c, m1 models)**. As such, it is difficult to correlate overall drug affinity to WT, inactivation deficient (S620T S641T), and inactivation enhanced (S641A) models based on interface score alone. Overall, a combination of solvent accessibility, drug orientation, and drug conformation as well as drug-protein contacts play a role in determining GALigandDock-based ligand interface scores.

While docking scores may not correlate with drug affinity, nearly all top-scoring ligands (except dofetilide-0/1 in S641T m2) made contact to known key drug binding residues Y652 and F656, corroborating experimental evidence (Fernandez et al., 2004; Mitcheson et al., 2000; Mitcheson and Perry, 2003; Perry et al., 2010; Sanguinetti and Mitcheson, 2005). An interesting aspect of this is that nearly all π-stacking interactions were made with Y652 (exceptions again arising in the S641A models with fenestrations), with F656 providing general hydrophobic interactions to the ligand. This strongly corroborates experimental findings indicating that mutation of Y652 to other aromatic residues is necessary for maintaining drug affinity, whereas bulky aliphatic groups were enough to approximate van der Waals interactions of F656 (Fernandez et al., 2004). However, residues at the bottom of the SF known to be critical for drug binding, T623, S624 and V625, were only accessible to some top-scoring models in the S641A model with fenestrations, suggesting an expanded pore cavity may be necessary for access.

### A potential role of R582

Lastly, we propose a molecular mechanism for entering the hERG inactivated state via a displacement of R582, which surrounds the extracellular gate in hERG m2 models, but sits atop the S6 segment, sandwiched between the S5P linker and the S5 helices in the relaxed hERG m1 models. A proposed mechanism outlining the order of events for hERG inactivation based on ϕ-value analysis compares the kinetic and thermodynamic relationship of the transition between two stable states (in this case open conducting and inactivated) using point mutations to identify the relative timing of domain motions and associated free energy changes in this transition (Wang et al., 1997). In the hERG channel, it was found that inactivation involves sequential conformational rearrangement of multiple segments, not just around the conducting pore (Perry et al., 2015), and our results place R582 in the hERG m2 models (which we propose are closer to an open state) near the path of the first step of inactivation (outgoing potassium ions), and near the subsequent second and third steps (S5 and S5P perturbation) in the hERG m1 models, including the S641A model with fenestrations, which we propose is close to an inactivated state. While clearly in need of experimental validation, our simulations enable us to explore structural molecular mechanisms of hERG inactivation.

## Discussion

The Rosetta based WT and mutant hERG models presented here provide structural insights into hERG inactivation and associated drug interactions. Our structural hERG models associate inactivation with an inward movement of F627 towards the ion conduction pathway. This movement is associated with loss of a hydrogen bond between S641 (in WT hERG) and N629, with a subsequent increase in hydrogen bonding between Y616 and N629 resulting in a shift of F627 into the conduction path. Additionally, a maximum lateral shift of F627 is associated with a loss of hydrogen bonding between the hydroxyl oxygen of S620 and the backbone amide hydrogen of F627, causing significant constriction of the SF backbone, which could impede ion flow as described for C-type inactivation (Cuello et al., 2010b; Li et al., 2017; Li et al., 2018; Li et al., 2021).

We have shown a potential correlation between the hERG S641A inactivation-enhancing mutation and the presence of four lateral fenestrations in the pore. This structural feature is accompanied by an increase in the conformational freedom of S5 and S6 pore-domain-forming segments, and the subsequent increase in pore radius. Each of four fenestration regions is of a strongly hydrophobic nature and is situated between residues F557, Y652 and F656, sitting below the “hydrophobic patches” behind the SF of the hERG WT cryoEM structure (Wang and MacKinnon, 2017). The hydrophobic pockets of the fenestrations in the S641A model may provide alternative binding mechanisms for hERG-blocking drugs during inactivation channel gating transition.

hERG S620T and S641T non-inactivating mutants demonstrate potential interactions that may block the movement of F627 into the ion conduction axis thus attenuating inactivation. Orientation of the threonine at the 620 position is identical across all m1 and m2 models, suggesting a stable conformation of the mutant. The orientation of threonine at the 641 position differs between m1 and m2 models, with the latter showing a higher probability for inactivation block by stabilizing F627 in the open conformation through hydrophobic interactions. Our findings for both S641A and S620T are corroborated by recent molecular dynamics simulations, suggesting energetically plausible mechanisms for inactivation or its block (Li et al., 2021; Miranda et al., 2020). An interesting aspect of the cryoEM hERG channel structure upon which our models are based is that it has a F627 backbone interchain carbonyl oxygen (S1 K^+^ binding position) distance of 6.3 Å, with F627 positioned away from the conduction path (Wang and MacKinnon, 2017). The authors suggest that the structure may be in a possible inactivated state when compared to a non-inactivating S631A mutant with F627 shifted slightly into the conduction path. However, a more recent cryoEM structure of a proposed inactivated state of hERG also shows F627 pointed away from the conduction path, but with a nearly 2 Å increase between carbonyl oxygens in the S1 position; inactivation in this case being the loss of K^+^ ion coordination, not necessarily the orientation of F627 (Asai et al., 2021). In comparison, our m1 and m2 cryoEM-refined starting models have S1 diameters of 7.2 Å; more open than their archetype hERG structure, but clearly more constricted than the newly proposed inactivated structure. Thus, the use of a proposed SF collapse alone, as a sign of C-type inactivation, could not be implied from our models in large part due to the absence of explicit water molecules and K^+^ ions that play a crucial role in forming the structural features associated with the channel conduction and inactivation. Additionally, residue packing optimization during the Rosetta Relax protocol naturally causes repacking of sidechains in the protein structure, resulting in significant conformational changes of the SF in all models. We therefore combine SF collapse with a shift of F627 into the conduction path and a loss of hydrogen bonding between S620 and F627 – elements all found in our S641A m1 models – as a potential mechanism of C-type inactivation.

Computational models of hERG potassium channel provide structural insights into an inactivated state and associated drug interactions. Our computational approach will be useful to study ion channel modulation by small molecules.

## Acknowledgements

We thank Drs. Heike Wulff and Jie Zheng and members of the V.Y.-Y., C.E.C., I.V., and J.T.S. laboratories for helpful discussions.

## Authorship Contributions

Participated in research design: Jan Maly, Aiyana M. Emigh, Kevin R. DeMarco, Kazuharu Furutani, Jon T. Sack, Colleen E. Clancy, Igor Vorobyov, and Vladimir Yarov-Yarovoy

Conducted experiments: Jan Maly, Aiyana M. Emigh, and Kevin R. DeMarco. Contributed new reagents or analytic tools: Jan Maly, Aiyana M. Emigh, and Kevin R. DeMarco.

Performed data analysis: Jan Maly, Igor Vorobyov, and Vladimir Yarov-Yarovoy

Wrote or contributed to the writing of the manuscript: Jan Maly, Aiyana M. Emigh, Kevin

R. DeMarco, Kazuharu Furutani, Jon T. Sack, Colleen E. Clancy, Igor Vorobyov, and Vladimir Yarov-Yarovoy

## A list of nonstandard abbreviations used in the paper

AP: action potential
cryoEM: cryo-electron microscopy
ECG: electrocardiogram
hERG: human Ether-à-go-go-Related Gene
IKr: rapid delayed rectifier K^+^ current
Kv: voltage-gated K^+^ channel
LQTS: Long QT syndrome
P-helix: pore helix
PLIP: Protein-Ligand Interaction
server SF: selectivity filter
TEA: tetraethylammonium
VSD: voltage-sensing domain
WT: wild-type

## Footnotes

This work was supported by National Heart, Lung, and Blood Institute [Grants 5R01HL128537, 2R01HL128537, 1R01HL152681, 5U01HL126273]; NIH Common Fund [Grants 1OT2OD026580-01, 3OT2OD026580-01]; American Heart Association Predoctoral Fellowship [Grant 16PRE27260295]; American Heart Association Career Development Award [Grant 19CDA34770101]; and Department of Physiology and Membrane Biology Research Partnership Fund.

## Supplemental Material Methods

### S1 Text Rosetta movemap files and generalized command lines

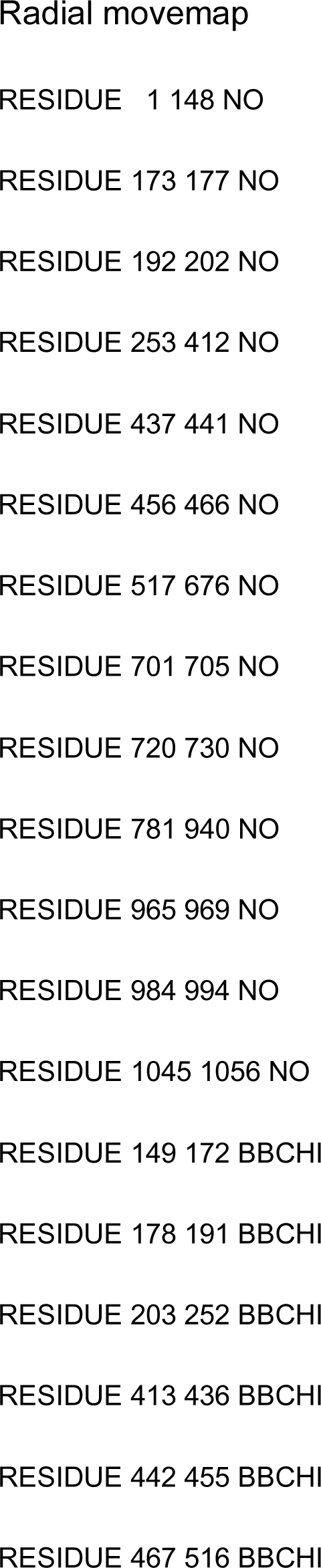
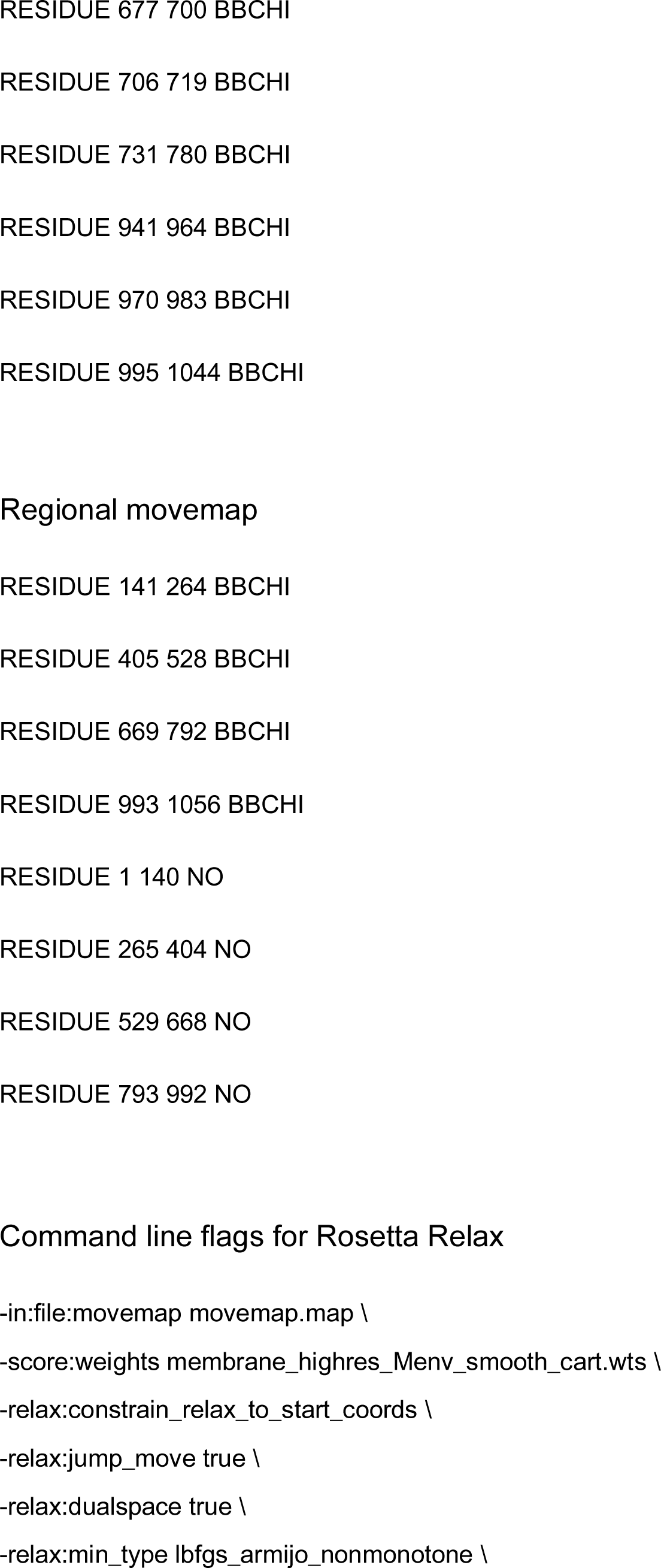
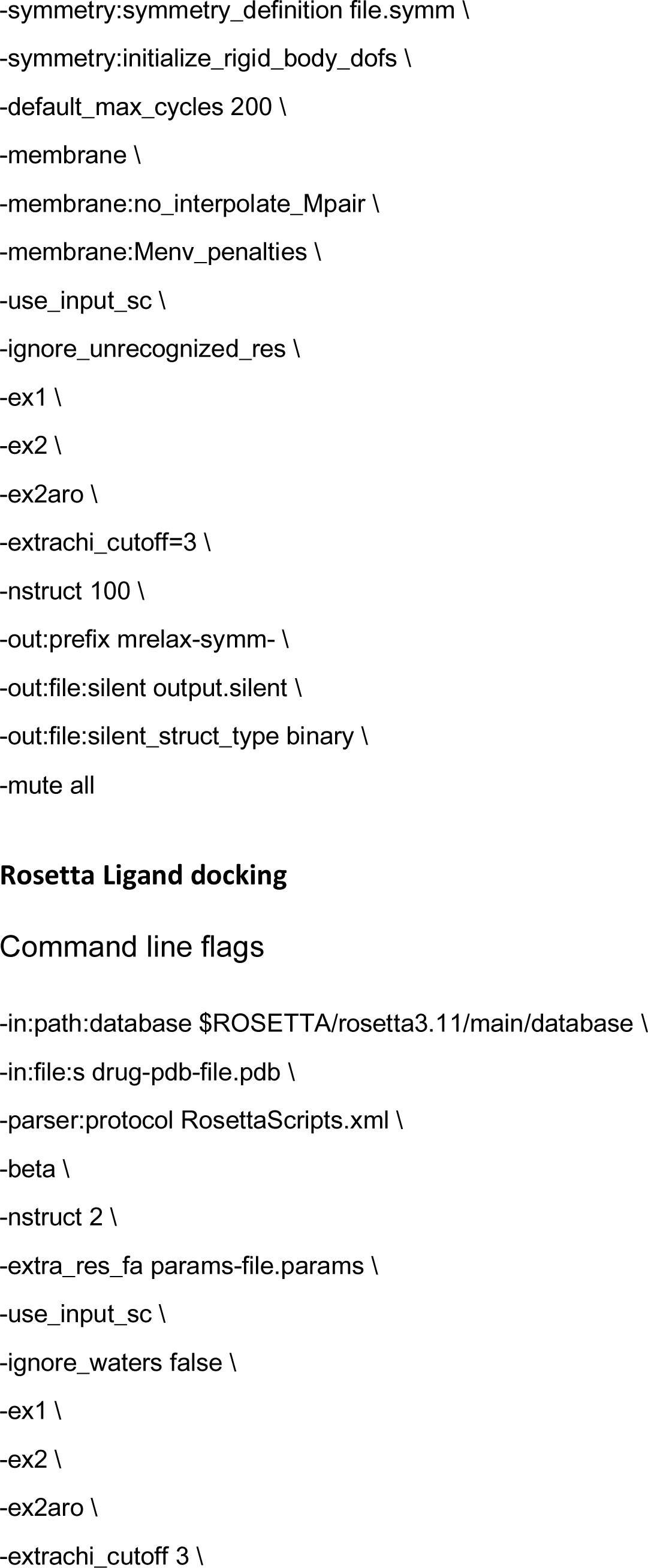
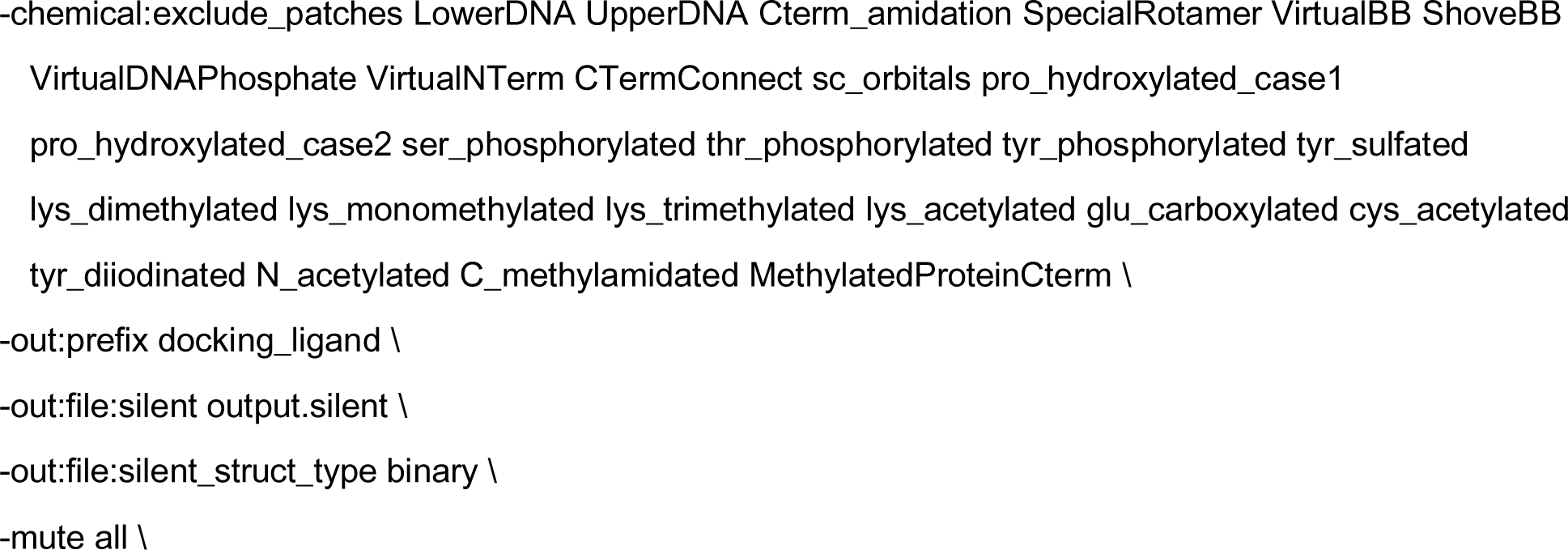

### S1 XML Script RosettaScripts for GALigand Dock

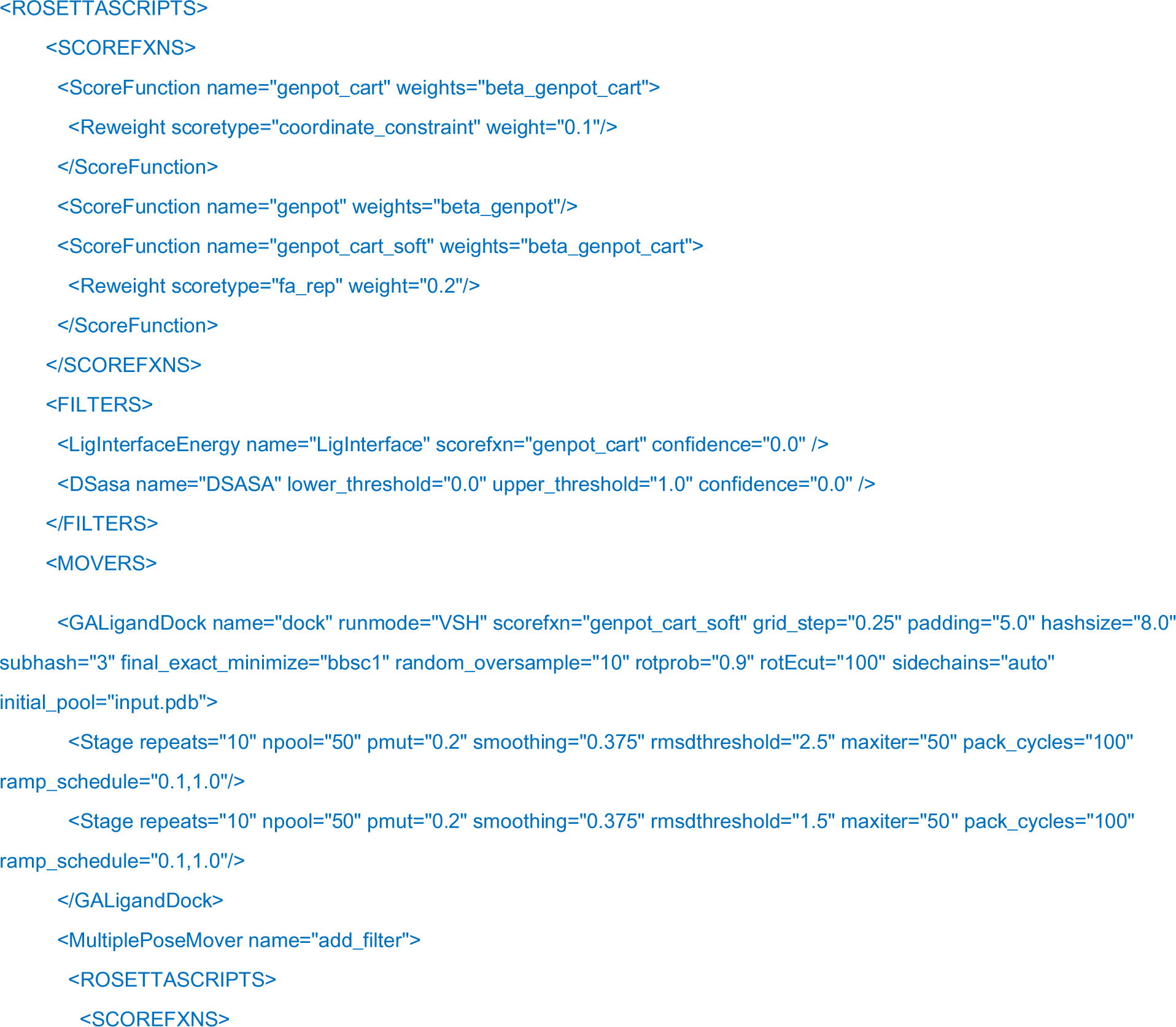
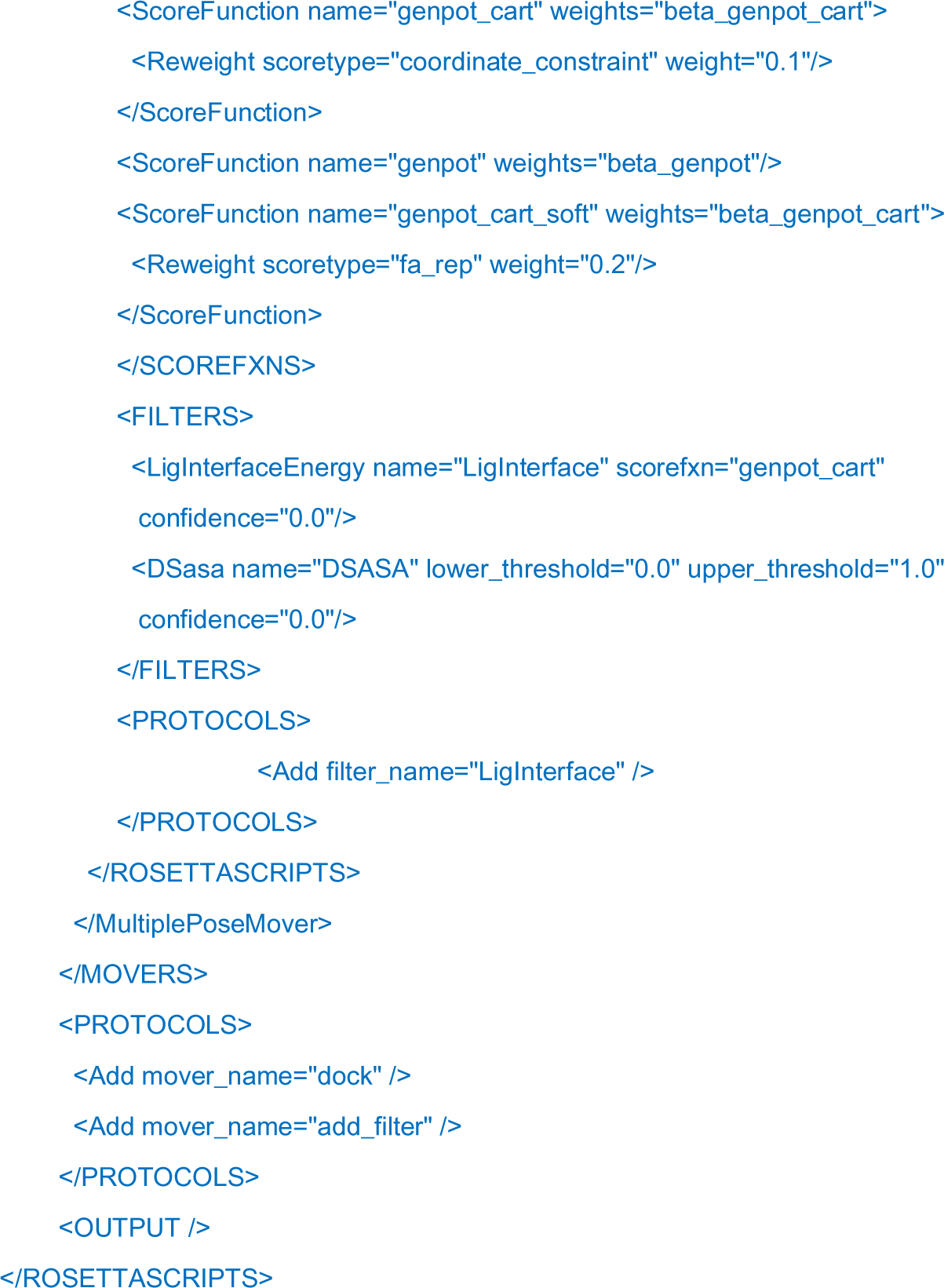

---

**Supplemental Fig. 1:**
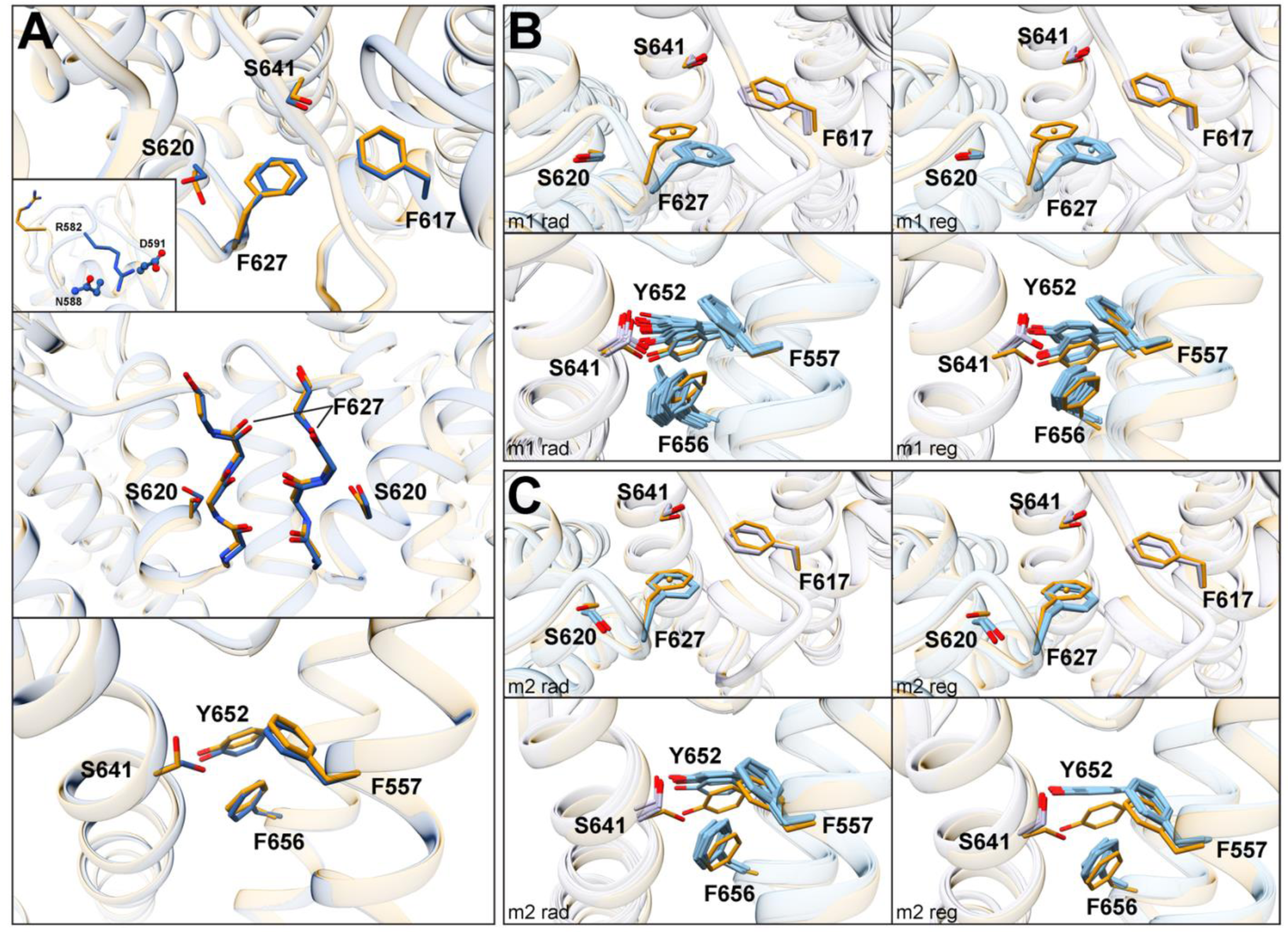
Comparison of input and top 20 relaxed wild type (WT) models. **A**) Overlay of model 1 (m1, orange stick) and model 2 (m2, blue stick) WT input models showing a top-view of F627 and surrounding residues (top), orientation of R582 on the S5P turret helix (top, inset), side-view of the SF backbone (middle), and backside-view of the hydrophobic fenestration region under the SF along S5 and S6 segments (bottom). **B**) Top 20 relaxed WT m1 radial (left panel) and regional (right panel) models showing conformational variation in F627 and surrounding residues (top) and fenestration region (bottom). Input structures are in orange stick, with all other residues colored according to specific domains as in **Figure 1**. **C**) Same as **B** but for WT m2 models.

**Supplemental Fig. 2:**
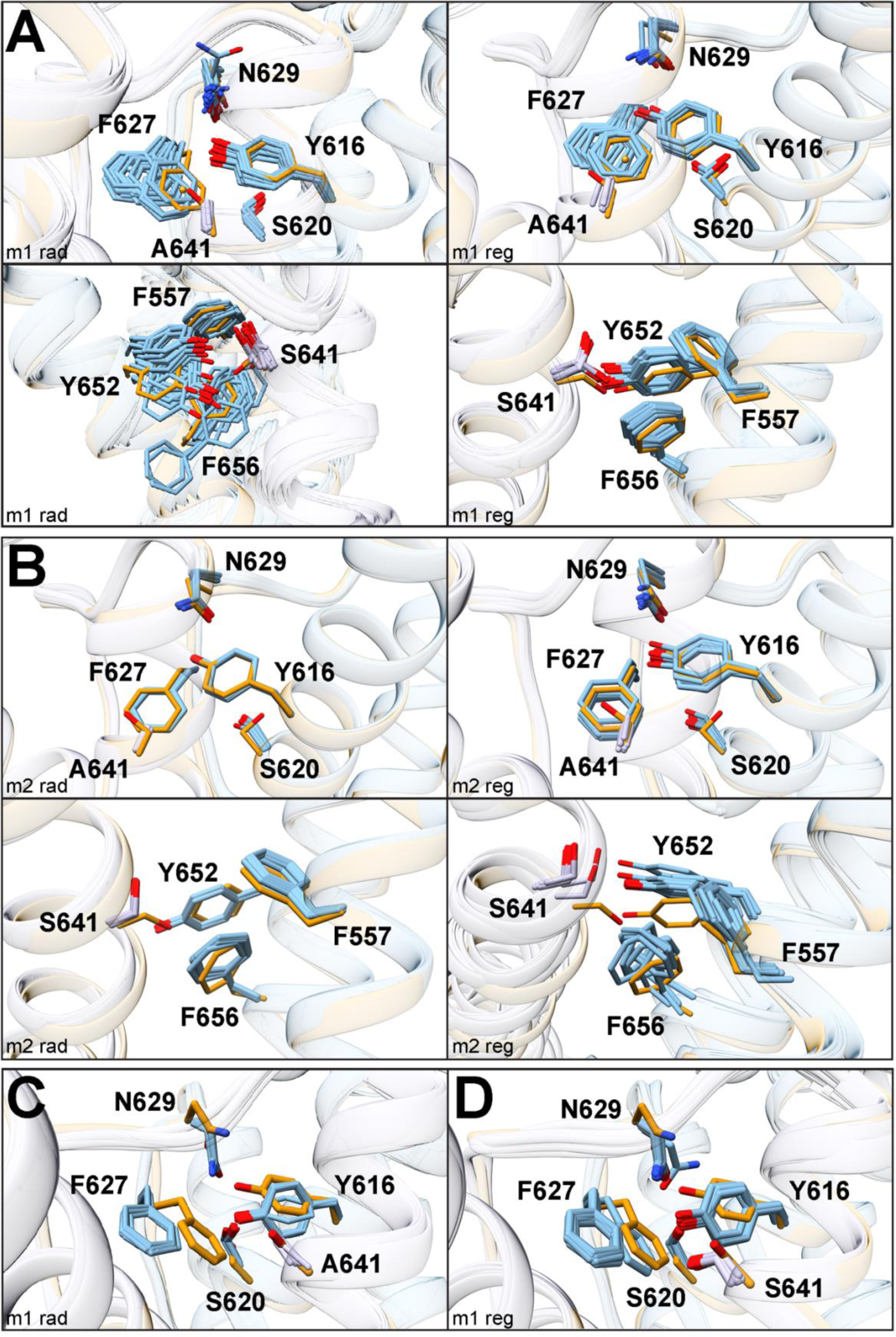
Comparison of top 20 relaxed S641A mutant models. **A**) Overlay of S641A m1 radial (left panels) and regional (right panels) models showing the SF (top) and fenestration (bottom) regions. Residue colors as in **Supplemental Fig. 1**. **B**) Same as **A** but for S641A m2 models. **C**) SF region of WT m1 radial models with a constrained S620-N629 hydrogen bond. **D**) SF region of S641A m1 regional models with a constrained S620-N629 hydrogen bond.

**Supplemental Fig. 3:**
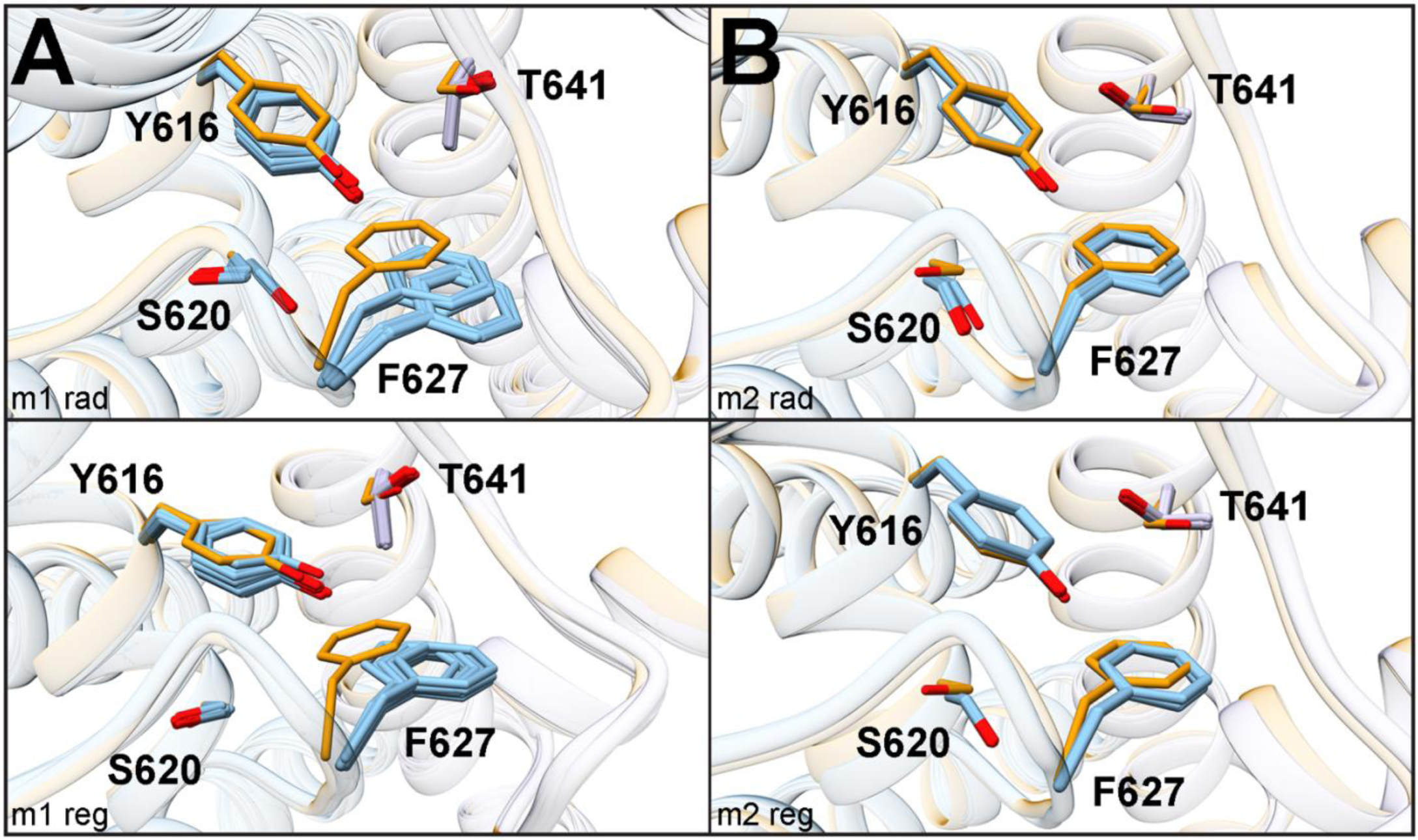
Comparison of SF region only for top 20 relaxed S641T m1 (**A**) and m2 (**B**) models for both radial (top) and regional (bottom) models. Residue colors as in **Supplemental Fig. 1**.

**Supplemental Fig. 4:**
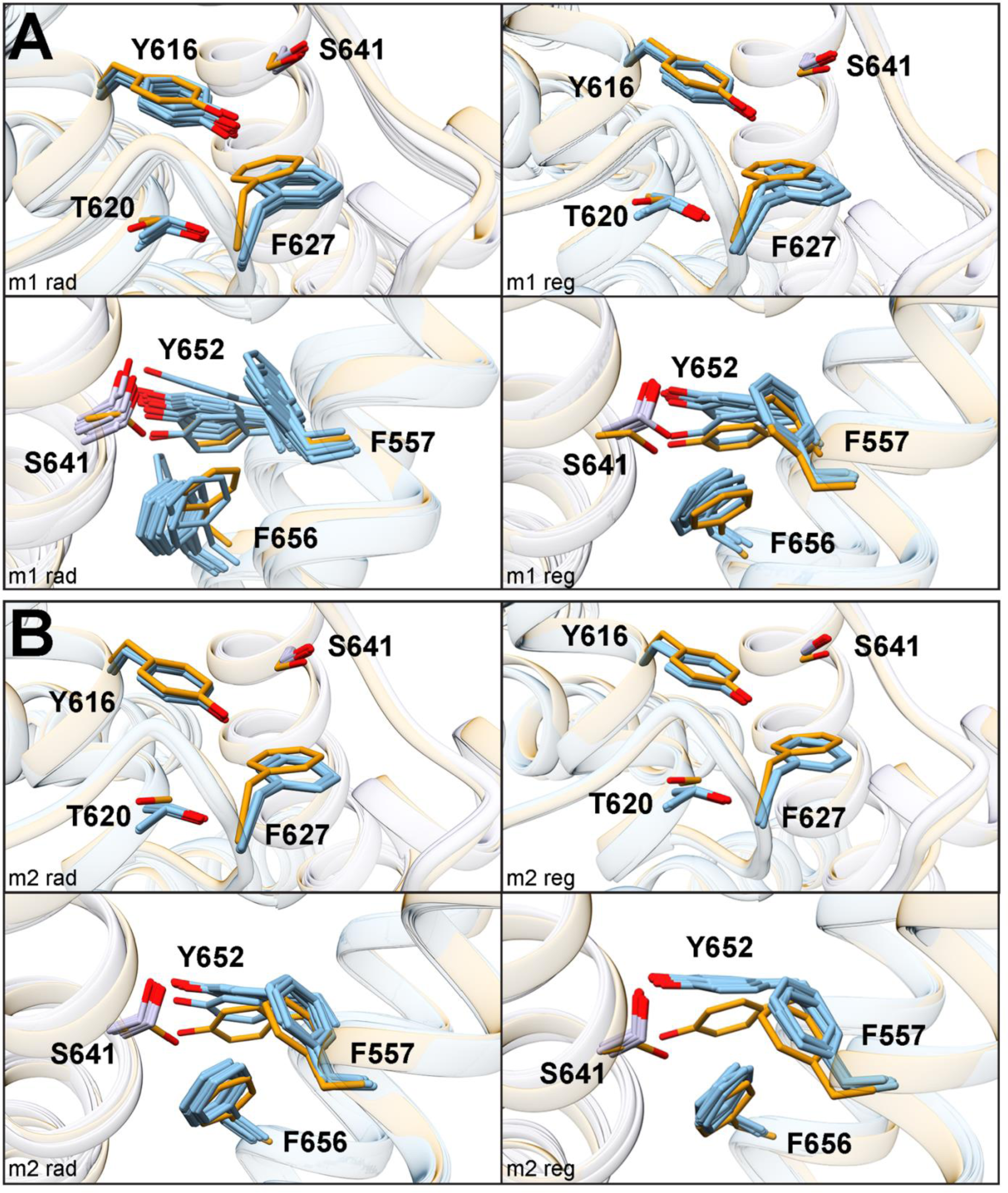
Comparison of top 20 relaxed S620T mutant models. **A**) Overlay of S620T m1 radial (left panels) and regional (right panels) models showing the SF (top) and fenestration (bottom) regions. Residue colors as in **Supplemental Fig. 1**. **B**) Same as **A** but for S620T m2 models.

